# Color, size, shape: The drivers of floral variation in *Hesperis matronalis* (Dame’s Rocket)

**DOI:** 10.1101/2025.10.16.682931

**Authors:** Stephen E. Johnson, Joanna L. Rifkin, Stephen I. Wright, Regina S. Baucom

## Abstract

Evolutionary biologists have long been intrigued by the factors that sustain genetic and phenotypic variation within and among natural populations. Polymorphisms underlying components of floral display – such as floral color, size, and shape – are uniquely of interest since variation in these traits impact pollinator attraction, rates of visitation, and pollinator efficiency, which ultimately influence patterns of plant reproduction and therefore fitness. We leverage existing floral variation present within and among populations of *Hesperis matronalis* (Dame’s Rocket) to disentangle the relative influence of natural selection and genetic drift in shaping floral trait variation. We employ a multi-tiered approach: we determine if variation in floral traits (color, size, and petal shape) is influenced by geography or environmental variables such as temperature and precipitation, we evaluate whether selection underlies trait variation by comparing phenotypic divergence (P_ST_) with neutral genetic structure (F_ST_), and we perform a Lande-Arnold selection analysis to explore the relationship between fitness and floral trait variation within natural populations. We find that selection underlies the divergence of floral color, floral size, and petal width among *H. matronalis* populations, with P_ST_ > F_ST_ for each trait. We find no indication, however, that variation in size in this species is influenced by the environment, but some evidence that variation in floral color and petal shape may be influenced by temperature. Finally, selection analyses of contemporary populations indicate divergent selection affecting combinations of color, petal shape, and plant size. These results suggest that the variation in floral shape in this species may be maintained due to environmental pressures, whereas floral color is influenced by pollinator visibility and the presence of different pollinator groups.

## Introduction

The presence of trait variation within populations has long fascinated evolutionary biologists. Such variation may be neutral, and subject to random forces, or under selection but transient, such that not enough time has passed for fixation of a favored type. Alternatively, within-population variation is often thought to be maintained through forms of balancing selection (Hedrick 2007; Delph and Kelly 2014). Distinguishing among these possibilities remains a central challenge in evolutionary biology.

In plants, striking variation in floral traits—such as color, size, and shape—is most often attributed to pollinator-mediated selection and the evolution of correlated traits leading to pollinator syndromes (Fenster et al. 2004; Rausher 2008; Dellinger 2020). However, within-population variation in floral traits, especially floral color, is well documented across plant species (Kay 1978,Galen 1999b), and a variety of hypotheses have been proposed to explain its presence. For example, in wild radish, pollinators preferred yellow or white flowers over pink or bronze morphs (Stanton et al. 1989), while herbivores also preferred and performed better on yellow/white morphs (Irwin et al. 2003), creating antagonistic selection on floral color. In *Ipomoea*, an allele conferring white flowers shows a transmission advantage when rare (Subramaniam and Rausher 2000), leading to the maintenance of variation *via* balancing selection. Allelic variants associated with floral color have also been shown to influence abiotic stress responses, including drought and heat tolerance (Coberly and Rausher 2003a; Strauss and Whittall 2006; Schemske and Bierzychudek 2007; Vaidya et al. 2018), raising the possibility that pleiotropic effects of allelic variation on different traits maintains variation via antagonistic or environmentally variable selection.

By contrast, although floral size and shape also vary within species, these traits have been far less frequently examined in the context of within-population variation compared to floral color. This gap is striking given that size and shape strongly influence pollinator interactions, affecting both attractiveness to visitors and the efficiency of pollen transfer (Campbell et al. 1996; Parachnowitsch and Kessler 2010; Caruso et al. 2019). Although studies are more limited, variation in size and shape, like color, may be influenced not only by pollinator-mediated selection but also by non-pollinator biotic and abiotic pressures (Strauss and Whittall 2006). For example, in *Polemonium viscosum*, larger corollas are preferred by pollinating bumblebees, but also by non-pollinating ants that damage wider-flaring corollas, generating antagonistic selection (Galen 1999a,b). In the same species, smaller corollas are favored at higher altitudes under drought stress (Galen et al. 1999), showing that floral size and shape, like color, can be subject to multiple and often opposing selective pressures. Together, these examples highlight the potential for diverse agents of selection to act on floral morphology, yet the extent to which such pressures maintain within-population variation in size and shape remains far less understood than for floral color.

Many studies have identified selective pressures on floral traits and have explored the ecological and genetic factors maintaining floral polymorphisms (e.g., (Gómez 2003; Strauss and Whittall 2006; Rausher 2008; Caruso et al. 2019). However, most focus on selection acting on individual traits or trait pairs, leaving open how suites of floral and vegetative traits jointly influence plant fitness. Understanding these relationships is critical because selection rarely acts on traits in isolation—nonlinear and correlational selection can shape adaptive trait combinations and maintain variation within populations (Conner and Hartl 2004). Disruptive or correlational selection among traits such as flower size, display, and architecture may therefore play a key but underexplored role in maintaining floral diversity. Moreover, with a few notable exceptions (e.g. (Streisfeld and Kohn 2005; Schemske and Bierzychudek 2007; Berardi et al. 2016; Milano et al. 2016), most studies assessing the evolutionary forces shaping floral variation have not directly ruled out the influence of genetic drift by contrasting phenotypic differentiation in traits to neutral genetic differentiation (i.e. F_ST_/Q_ST_ comparisons).

The short-lived perennial or biennial plant *Hesperis matronalis* (L), commonly known as Dame’s Rocket, provides a particularly good model for examining the maintenance of within-population variation. One of its most distinctive features is a striking floral color polymorphism, with populations containing individuals that produce white, pale (a light lilac color), pink, or purple flowers (Mitchell and Ankeny 2001; Majetic et al. 2007; Francis et al. 2009); Figure 1 A-D). An unpublished 40-year study of *H. matronalis* populations within a deciduous riparian woodland showed that populations were generally stable and that variation in floral color was maintained during this time (discussed in (Mitchell and Ankeny 2001; Majetic et al. 2007; Francis et al. 2009)), suggesting that balancing selection may be acting on floral color variation in this species. However, despite long-standing interest in this floral color polymorphism among plant biologists, the factors acting to maintain it have not been formally studied. Equally distinctive, though less studied, *H. matronalis* also exhibits striking within-population variation in floral size and petal shape (Figure 2). Across individuals, petal widths range from very broad and rounded to thin and dissected in appearance (R.S. Baucom, personal observation; Figure 2), and floral size also varies considerably among individuals (Figure 2). Both small- and large-flowered plants display either broad or narrow petals (Figure 2). Together, the presence of striking variation in color, size, and petal shape makes *H. matronalis* an ideal system for testing the evolutionary processes that maintain floral trait diversity within populations.

**Figure 1.**
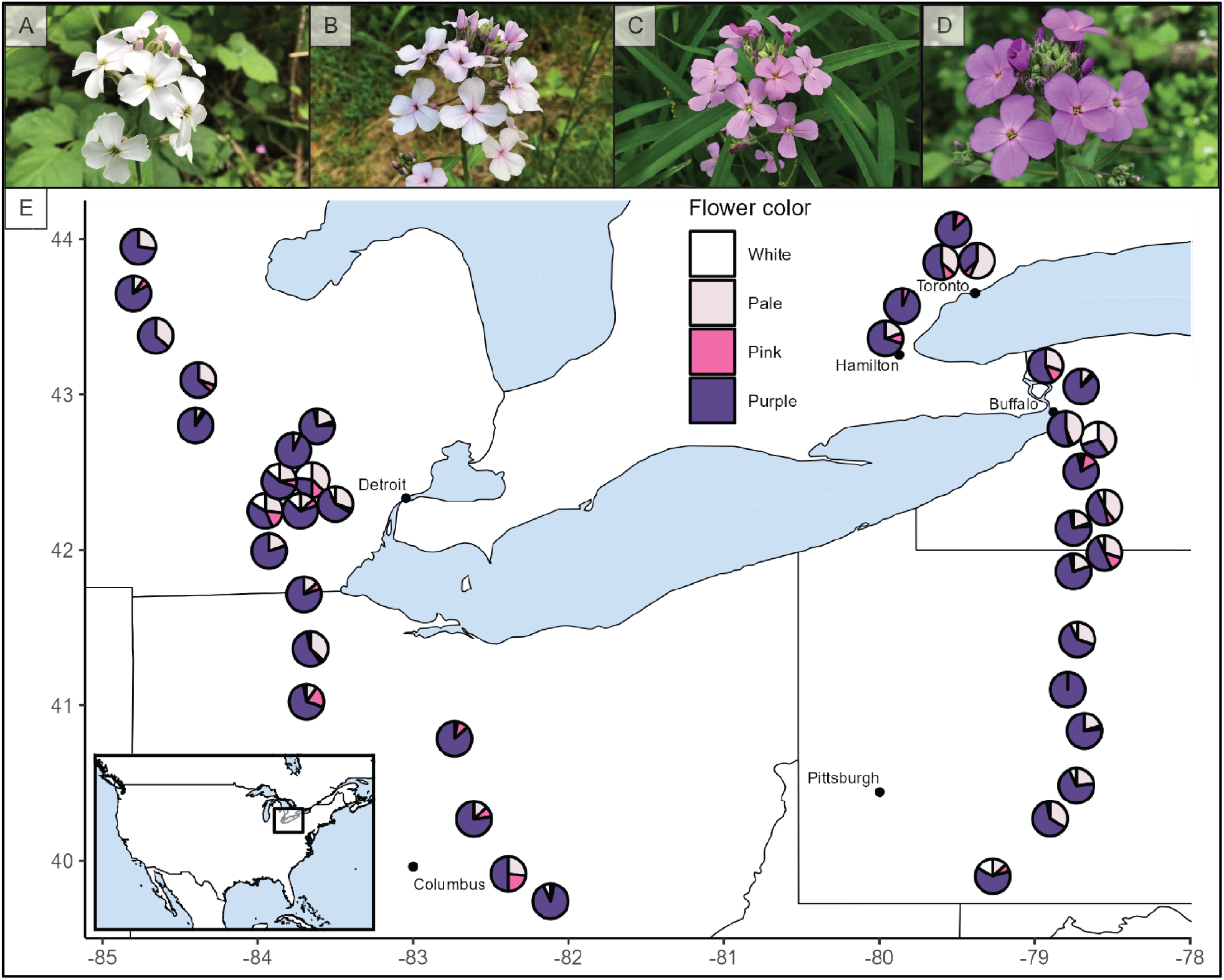
Images of *Hesperis matronalis* individuals and a map showing sampling locations from the United States and Canada in 2023. A-D) Images of representative white, pale, pink, and purple flowering individuals, respectively. E) A map showing the approximate locations of each sampling site indicating the proportion of individuals with white, pale, pink, or purple flowers, based on a visual assessment of a random subset of individuals within each population. A corresponding map with precise locations and population names is provided as Figure S1. Detailed population characteristics are available in Table S1.

**Figure 2.**
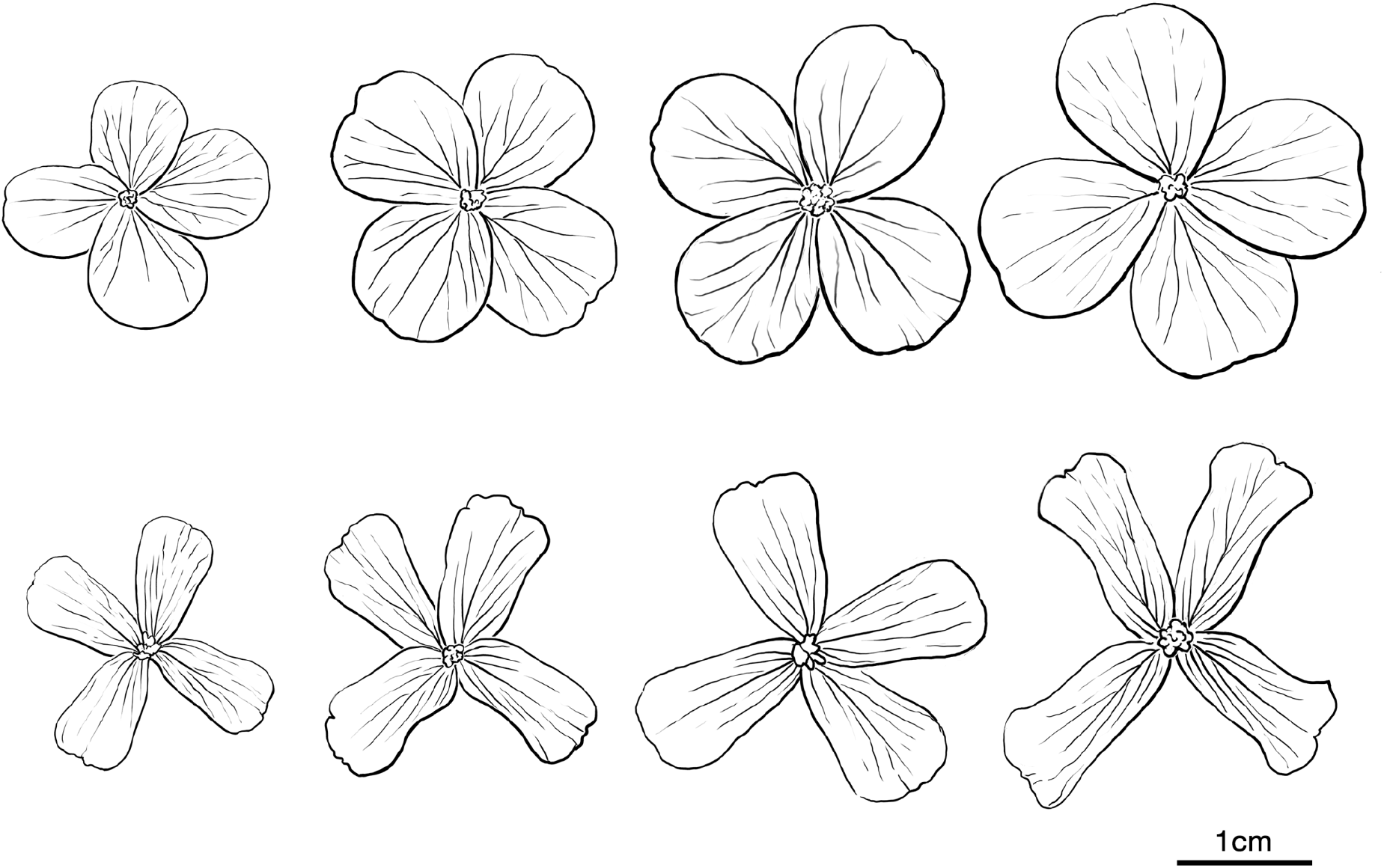
Line drawings by John Megahan (University of Michigan) illustrating the range of variation in floral size and petal width observed in *Hesperis matronalis*. Drawings were based on photographs of flowers collected from natural populations along both the Midwest and East transects. Variation is shown for overall flower size as well as petal width, which spans from broad and rounded to narrow and dissected morphologies. Scale bar = 1 cm.

To test whether floral variation in *Hesperis matronalis* is shaped by natural selection and potentially maintained by balancing selection, we examined flower color, size, and petal shape (measured as petal width) across replicated natural populations in North America. We first assessed whether floral traits varied clinally with geography or environment and quantified among-population differentiation. We then compared phenotypic divergence (P_ST_) with neutral genetic divergence (F_ST_) to test whether population differences exceeded neutral expectations. Finally, we performed Lande–Arnold selection analyses among populations to quantify the form and strength of selection acting on floral traits. This multi-tiered, field-based approach examines both landscape-level patterns of divergence with direct estimates of selection, providing a comprehensive framework for evaluating the evolutionary forces maintaining floral variation.

## Methods

### Study system

*Hesperis matronalis* (Dame’s rocket, Brassicaceae) is an herbaceous biennial native to Eurasia (Francis et al. 2009) believed to have been introduced to North America in the late 17th century (Al Gleason and Cronquist 1964). It was first reported as a garden escapee in the 19^th^ century and is now widespread across the East and Midwest United States and southern Ontario (Francis et al. 2009). North America has diploid and polyploid populations (Francis et al. 2009), with evidence suggesting the polyploid is a segmental allotetraploid (Gohil and Raina 1987). *H. matronalis* is hermaphroditic and self-compatible, with fewer seeds produced by selfed than outcrossed fruits (Mitchell and Ankeny 2001, Majetic 2008a). As in most members of the Brassicaceae, flowers of this species are radially symmetrical with four petals of the same size arranged in a cross shape (Nikolov 2019). Floral color variation in *H. matronalis* is due to changes in the anthocyanin pathway (Majetic 2008b), although the exact causal loci have not yet been identified in this species. Prior work shows that field-collected maternal lines, when emasculated and self-pollinated, produce individuals that typically exhibit the same floral color as the maternal parent (*i.e.*, white or purple flowered maternal plants most often produced white or purple flowered offspring, (Majetic 2008b)), suggesting that floral color in this species is primarily under genetic control. Species including bees, syrphid flies, and moths pollinate *H. matronalis* (Mitchell and Ankeny 2001, Majetic 2008a) with one study reporting no pollinator preference for flower color in Pennsylvania (Majetic et al. 2009).

### Population sampling for floral size, shape, & color

In May–June 2022, we sampled 20 *Hesperis matronalis* populations along a latitudinal transect from southern Ohio to central Michigan (“Midwest transect”; Fig. 1). At each site we recorded latitude, longitude, and altitude (Google Earth). We tagged 30–50 individuals ≥1 m apart, noting stem height, number of stems, and leaves (five bins: 0–9, 10–24, 25–49, 50–99, 100+). Leaf tissue from 10 plants per site was dried in silica for DNA extraction and ddRAD sequencing. For each tagged plant we scored flower color (purple, pink, pale, white) with a color chart and collected three newly opened flowers to assess morphology. Within 72 hr, flowers were photographed flat against construction paper with a scale, then traced in ImageJ to estimate floral area (cm^2^) and petal width (cm, at the broadest point). After senescence we collected siliques from each tagged plant; silique number, which strongly correlated with seed mass (N = 155, r = 0.85, p < 0.001), was used as a fitness proxy.

In May–June 2023, we resampled the 20 Midwest sites and added 20 new populations spanning Pennsylvania to Ontario (“East transect”; Table S1). The same sampling procedures were followed, but floral color was also measured quantitatively using a Flame Miniature spectrometer (Ocean Insight FLAME-S-UV-VIS-ES) in addition to qualitative scoring. For ∼30 plants per site, we measured standardized reflectance spectra (300–750 nm) from three flowers per individual after first calibrating the spectrometer with light and dark standards. Spectral curves showed peaks at 450 and 690 nm and a valley at 560 nm (Fig. 2A). Following Frey ((Frey 2004); (Frey 2007; Del Valle et al. 2018)) and Del Valle et al. (Frey 2007; Del Valle et al. 2018), we defined “depth” as average reflectance at 450 and 690 nm minus reflectance at 560 nm.

We analyzed color in two ways. First, we fit a generalized linear model (R v.4.2.3, glm) testing whether qualitative classes predicted spectral depth, using the following model:

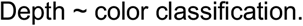

A Gamma distribution with log link provided best fit, and significance was assessed with Wald Type II χ^2^ tests. Post-hoc Tukey HSD contrasts were implemented with the emmeans package (Russell n.d.). Second, a PCA of depth, saturation, brightness, and reflectance at 450 nm, overlaid with qualitative scores, showed a strong separation between “light” (white, pale) and “dark” (pink, purple) morphs, but overlap between pink–purple and pale–white classes.

As in 2022, three leaves from 10 randomly chosen plants per East transect site were collected for DNA extraction and RADseq.

### Assembling abiotic environmental variables for observed populations

We evaluated environmental influences on floral traits (color depth, area, petal width) using 2023 data from both transects and 2022 floral size/petal width data from the Midwest transect. We examined both historical and contemporary environmental variables since we reasoned that historical environmental estimates likely reflect conditions under which populations adapted, and contemporary estimates capture more recent temperature and precipitation patterns to which populations are exposed. We focused on temperature and precipitation, as prior studies suggest both can influence floral color (Schemske and Bierzychudek 2001,Coberly and Rausher 2003b). For historical environments, we extracted 19 temperature- and precipitation-related bioclimatic variables (“bioclim”) from WorldClim (http://www.worldclim.org/) and reduced dimensionality via PCA (prcomp, R Core Team 2023). One PCA included all sites, and a second included only Midwest sites because trait data differed by transect. Principal components explaining ≥20% of variance were retained for analysis.

For analysis of contemporary environments, we examined temperature and precipitation during the species’ reproductive stage (*i.e.*, May-June) from the year of population sampling. We elected to perform analyses using June temperature and precipitation values since *Hesperis matronalis* typically flowers and sets seed in a short period of time between early May and end of June in the areas in which populations were sampled. Preliminary analysis indicated June variables were most explanatory, so we used daily low and high temperatures and precipitation from the nearest NOAA weather station for U.S. sites (rnoaa v1.4.0; (Chamberlain and Hocking 2023)) and from Environment Canada (http://www.weather.gc.ca; accessed July 9, 2024) for Canadian sites. From daily data, we calculated mean June daily lows, mean June daily highs, and cumulative June precipitation. Absolute pairwise distances between populations were then calculated for each variable and composite PC.

### Phenotypic population structure

We tested whether floral and vegetative traits showed significant population structure, reasoning that such structure could reflect either natural selection or drift. For the Midwest transect (2022), we examined flower size (area), petal width, stem number, height, and leaf number; for 2023, we tested whether flower color depth varied within and between the Midwest and East transects. Floral size and petal width were averaged per individual. Trait distributions were checked for normality and transformed as needed (bestNormalize; (Peterson 2021b).

Vegetative traits and floral size traits from 2022 were analyzed using linear models of the form:

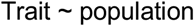

with population as a fixed effect; significance was assessed with Type II sums of squares (car package; (Fox and Weisberg 2018). For color depth, we fit generalized linear models in R v4.2.3 (glm, Gamma distribution, log link) with the model:

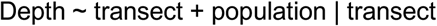

treating transect and nested population as fixed effects. Depth values were averaged per individual, and significance tested as above.

Because replicate flowers from the same plant appeared similar in size and width, we further tested whether variance was greater among individuals than among flowers within individuals. We fit GLMMs with glmmTMB (Brooks et al. 2017):

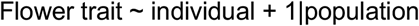

with individual as a fixed effect and population as a random effect. We specified lognormal error for floral area and tweedie error for petal width, with diagnostics evaluated using DHARMa (Hartig 2024).

### Neutral genetic markers for population differentiation

We extracted DNA from leaves using DNeasy Plant Mini Kits (Qiagen) and prepared four ddRAD libraries of 96 samples each following Peterson et al. (2012; protocol in Supplementary Material). Libraries were sequenced at the University of Michigan Advanced Genomics Core (151-PE, NovaSeq). Across the four libraries, 192 samples from 20 Midwest transect sites and 192 from 20 East transect sites were sequenced (N = 384 total, 8–10 individuals/site), yielding 1,293,713,823 raw read pairs (mean = 3,374,255 per sample; Table S3).

Fastq files were demultiplexed and adapter dimers removed with cutadapt (Martin 2011), using barcodes identified by FastQC (Andrews 2010). Reads were aligned to a reference genome (cite) with BWA-MEM (Li 2013), sorted with SAMtools (Danecek et al. 2021), annotated with read groups, and PCR duplicates removed using (Picard toolkit 2019). Ploidy was assessed with the nQuire “lrdmodel” function (Weib et al. 2018). After filtering, 548,404,618 read pairs remained (mean = 1,428,138/sample; Table S3).

SNPs were called with FreeBayes v1.3.6, a haplotype-based variant caller appropriate for tetraploid genomes (Garrison and Marth 2012), using conservative quality filters optimized for many samples and the ploidy -4 option. SNPs were retained if quality scores exceeded 20 and genotypes had ≥ 5× coverage (BCFtools; (Danecek et al. 2021). The VCF was iteratively filtered to keep SNPs present in > 90% of individuals and individuals with > 75% SNP calls. Because aligning tetraploid samples to a diploid reference can produce spurious fixed heterozygous calls from homoeologous alignment (that are not bona fide polymorphisms, but rather fixed differences between gene duplicates), we removed sites with allele frequency = 0.5 using SAMtools view, then excluded multi-allelic SNPs (full protocol in Supplementary Material).

Analyses were performed on all samples combined and separately by transect, yielding three VCFs: all individuals (N = 299,692 SNPs), East transect (N = 142,6049 SNPs), and Midwest transect (N = 133,583 SNPs). Results primarily focus on the combined dataset, with transect-specific outcomes noted when they differ.

### Analysis of neutral population structure and isolation by distance

We assessed genetic diversity, structure, and isolation by distance (IBD) in natural populations of *Hesperis matronalis* using R v4.2.3 ((R Core Team 2023) in RStudio v2023.03 (Posit team 2023). Expected and observed heterozygosity were estimated with adegenet v2.2.10 ((Jombart 2008; Jombart and Ahmed 2011). Global F_ST_ across all populations and within each transect was calculated using the “basic.stats” function in hierfstat v0.5-11 ((Goudet 2005), with 95% confidence intervals derived from jackknife standard errors. Pairwise F_ST_ among populations was calculated with “stamppF_ST_” in StAMPP v1.6.3 ((Pembleton et al. 2013), which implements Weir and Cockerham’s F_ST_; 95% CIs were obtained by bootstrapping.

To partition genomic variation, we conducted an AMOVA with “poppr.amova” in poppr v2.9.6 ((Kamvar et al. 2014), evaluating variance among transects, among populations within transects, and within populations. Departure from panmixia was tested with 999 randomizations using “randtest” in ade4 v1.7-22 ((Dray and Dufour 2007).

To test for IBD, we performed Mantel tests between pairwise geographic distances and pairwise F_ST_ values. Geographic distances were computed with “earth.dist” in fossil, and Mantel tests with 10^6^ permutations were implemented using “Mantel” in ecodist v2.1.3 ((Goslee and Urban 2007).

### Admixture

We used the conStruct R package v1.0.6 (Bradburd et al. 2018) to estimate the number of subpopulations (K). Cross-validation was performed with the “x.validation” function for spatial and nonspatial models, testing K = 1–10 with 10 permutations each. The K with the lowest mean cross-validation error was selected for further analysis. Spatial models incorporate geographic location, assuming genetically closer individuals are geographically proximate due to IBD, whereas nonspatial models ignore location to assess differentiation from other factors (e.g., environmental or ecological). For the best-supported K, we ran 1000 permutations of both spatial and nonspatial models and report mean cross-validation errors.

### Comparing P_ST_ and F_ST_ for evidence of selection

We compared phenotypic differentiation in floral traits (P_ST_) to neutral genetic differentiation (F_ST_) to assess whether population structure reflected selection or drift. P_ST_, conceptually similar to Q_ST_, is based on phenotypic rather than additive genetic variance (Merilä and Crnokrak 2001; Leinonen et al. 2008; Whitlock 2008). If P_ST_ exceeds F_ST_, especially under ongoing gene flow, divergence likely reflects diversifying selection. If P_ST_ is lower than F_ST_, stabilizing selection is implicated; equivalence of P_ST_ and F_ST_ indicates that phenotypic differentiation aligns with what would be expected from genetic drift alone.

We assessed differentiation in traits among populations (P_ST_) using the formula:

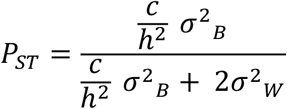

where c is the proportion of variance due to additive genetic effects among populations, h^2^ is within-population narrow-sense heritability, *σ*^2^_B_ is the among-population variance, and *σ*^2^_W_ the within-population variance. *σ*^2^_B_ was calculated as MS_B_ - MS_W_ / N, where MS_B_ is the between-population mean square, MS_W_ the within-population mean square (unbiased error variance), and N the number of individuals per site (Sokal and Rohlf 1995 p. 217). This approach has been widely applied to estimate P_ST_ or Q_ST_, including for non-normally distributed traits (Storz 2002; Brommer 2011).

P_ST_ and 95% CIs were calculated with the “Pst” function in Pstat v1.2 ((Silva and Silva 2018), modified to implement the above *σ*^2^_B_ formula. Because c and h^2^ are unknown, we evaluated P_ST_ across c/h^2^ ratios of 0.2, 0.5, and 1.0, representing conservative, intermediate, and upper estimates (Brommer 2011); (Morente-López et al. 2020). Ratios >1 are rarely used, as additive genetic variance among populations seldom exceeds that within populations (Brommer 2011).

We contrasted P_ST_ and F_ST_ for flower color (depth) using all populations, and for floral size (area), petal width, height, stem number, and leaf number using Midwest populations only. For each trait, global P_ST_ and F_ST_ were compared via 95% CI overlap. Pairwise P_ST_ – F_ST_ comparisons were conducted for flower color across all populations and for floral size and petal width within the Midwest. For most traits, c/h^2^ = 0.5 was assumed; for flower color, c/h^2^ = 1 was used, as color is likely controlled by one or two loci with additive effects and is nearly ubiquitous across populations ((Majetic et al. 2007).

### Clinal variation and geographically variable selection

We tested whether variation in floral and size traits of *Hesperis matronalis* was associated with environmental factors using two approaches:

1. Correlation with environmental and geographic variables

We calculated Pearson’s correlation coefficients between population means of plant phenotypes (flower color depth in 2023; flower size, petal width, stem number, height, and leaf number in Midwest populations in 2022) and environmental variables (WorldClim PC1, PC2, cumulative June precipitation, mean June daily low, and mean June daily high). Correlations were computed with Hmisc v3.12-2 (Harrell 2024), with sequential Holm–Bonferroni correction (Holm 1979); α = 0.05, using stats v3.0.1, (R Core Team 2023). Associations with geographic distance were tested using Mantel tests in ecodist v2.1.3 (Goslee and Urban 2007).

2. Multiple regression of distance matrices (MRM)

We used MRM to test whether environmental differences (WorldClim PC1, PC2, June precipitation, mean June low, and mean June high) explained phenotypic differentiation (P_ST_) among populations. Unlike partial Mantel tests, MRM accounts for correlations among predictors, partitions variance, and estimates effect sizes. Each model included pairwise F_ST_, geographic distance, and environmental differences, enabling assessment of their relative contributions to phenotypic divergence while controlling for IBD and population structure. Environmental differences were calculated as absolute between-population values.

All distance matrices (P_ST_, F_ST_, environment, and geographic distance) were standardized (mean = 0, SD = 1) using decostand in vegan v2.6-4 (Oksanen et al. 2017).

For each focal trait (flower color depth in both transects; floral area and petal width in the Midwest), we modeled phenotypic differentiation (P_ST_) with linear regressions to assess genetic and geographic effects. Four models were tested for each trait–environment combination:

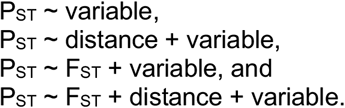

P_ST_ matrices were permuted 10^6^ times with the “MRM” function in ecodist (Goslee and Urban 2007) while explanatory matrices remained fixed. False discovery rate was controlled per trait using the Benjamini– Hochberg procedure (Benjamini and Hochberg 1995).

### Phenotypic selection analyses

We tested whether natural selection influenced variation in floral traits (flower color, area, petal width) and plant size traits (stem number, height) using Midwest collections from 2022, which included silique counts for 30–50 individuals per population. Flower size was measured in the field, and flower color scored qualitatively (white, pale, pink, purple). Unlike later analyses that used 2023 spectrometer data (“depth”), no spectrometer was available in 2022; therefore, field-based color categories were used. Because distinguishing pink from purple was difficult by eye, we binned individuals into “light” (white, pale) and “dark” (pink, purple) categories for selection analyses.

Phenotypic selection was quantified following Lande and Arnold (Lande and Arnold 1983). Relative fitness was calculated as individual silique number divided by the population mean. Phenotypic traits were standardized within populations (mean = 0, variance = 1). Linear (directional) selection gradients (β) were estimated from the model:

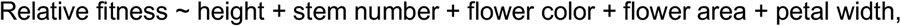

with flower color coded as light/dark. Leaf number was excluded due to strong correlation with height (r = 0.85, p < 0.001). Nonlinear selection gradients (γ) were estimated by doubling quadratic regression coefficients from models including linear, quadratic, and cross-product terms. Significant γ values indicate quadratic (variance) or correlational (covariance) selection. Three-way interactions among traits were also tested, with emphasis on interactions between plant traits and flower color (e.g., stem number × height × color). Four-way interactions were not included.

Because selection analyses revealed strong interactions between plant size traits (stem number, height) and flower color (see Results), we conducted linear and quadratic selection analyses separately for dark and light morphs. Linear selection gradients (β) were estimated from:

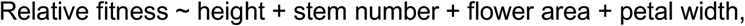

Nonlinear gradients (γ) were estimated from:

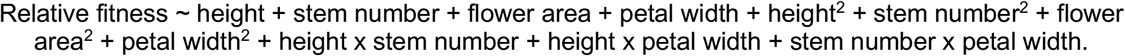

Fitness surfaces for significant correlational selection were visualized with thin-plate spline regression (Tps function, fields; (Furrer et al. 2010), with smoothing parameters chosen by minimizing generalized cross-validation scores.

To test whether trait values differed between color morphs, we ran mixed-model ANOVAs for fitness, flower size, petal width, stem number, and height:

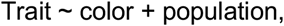

where color was a fixed effect and population a random effect (lme4, (Bates et al. 2011) . Trait distributions were evaluated for normality with bestNormalize (Peterson 2021a) and transformed as needed: asinh (fitness, height), log (stem number, petal width), or square root (flower area). Significance of color effects was assessed using Wald Type II Chi-square tests.

## Results

### *Flower traits and trait variation across natural populations of* Hesperis matronalis

Using spectrometry data sampled from natural populations in 2023, we validated our qualitative flower color classifications of white, pale, pink, and purple (Figure 3A,B). Spectral depth differed strongly among the four visually defined classes (χ^2^_3_ = 2448, P < 0.001), with post-hoc comparisons confirming significant pairwise differences in mean depth (mean ±se, white: 2.8 ± 0.4, pale: 10.1 ± 0.5, pink: 28.4 ± 0.9, and purple: 34.2 ± 0.3, Figure 3B). A PCA of reflectance traits (depth, saturation, reflectance curve height at 450 nm, brightness) indicated that while the four groups are largely distinct, some overlap occurs between pinks and purples and between whites and pales (Figure 3B, Figure S3). Overall, these analyses show that *H. matronalis* flower color variation forms four primary peaks in spectral space, with some intermediate phenotypes bridging classes.

**Figure 3.**
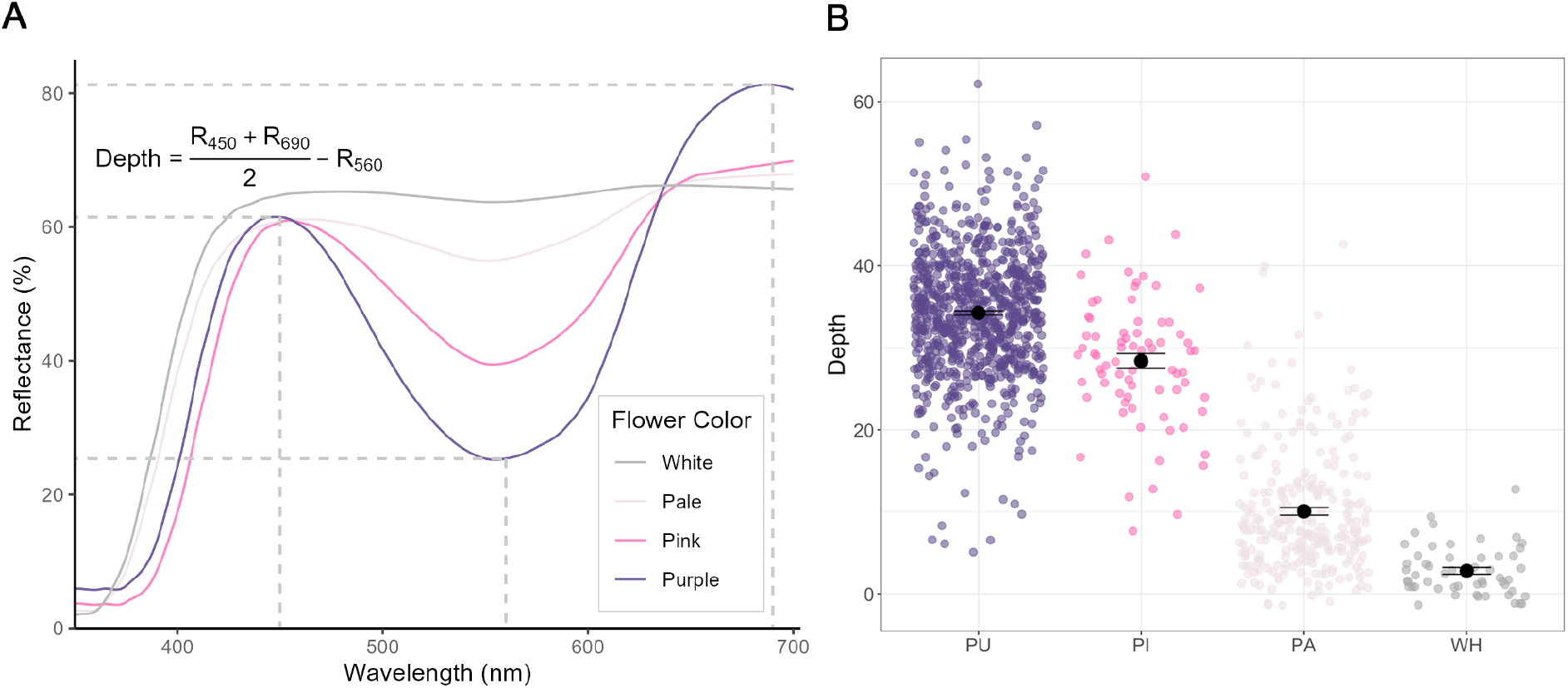
Floral color variation in *Hesperis matronalis*. A) Example reflectance curves from spectrographic analysis of the four main floral-color classes sampled across natural populations in 2023. The dashed grey lines on the purple curve indicate reflectance values at wavelengths used to calculate depth. Reflectance curves exhibiting depth values close to the species average for each of the four color classes (*i.e.* white, pale, pink, and purple) are presented for comparison. B) Values of depth from each individual (averaged from three flowers per plant) sampled in the field in 2023 and presented by floral color class determined in the field (PU = purple, PI = pink, PA = pale, WH = white). Black dots indicate the average (±se) depth value corresponding to each floral color class. On average, depth significantly differed between the field-based estimates of floral color, but the variation between classes suggests that *H. matronalis* exhibits four main peaks of floral color, with continuous color variation between peaks.

Flower size and petal width exhibited ample variation among individuals (Figure 2). The average flower size (area) across individuals ranged from 0.85–6.32 cm^2^, with a species mean of approximately 2.8 cm^2^. Petal width ranged from 0.37–1.38 cm, with a mean of ∼0.9 cm. Individual plants differed significantly for both traits (Table S4), consistent with our general observation that flowers from a given plant typically exhibit the same flower size and petal width. The range of values indicates substantial variation among individuals, particularly in flower area where the largest flowers were more than seven times the size of the smallest.

Populations also differed significantly for flower color, flower size, and petal width. Mean spectral depth values ranged from 13.3–38.7 across populations (χ^2^_39_ = 202.5, P < 0.001; Figure 4, Tables S5– S6), a difference of ∼190% between average values of the lightest- and darkest-flowered populations. Although populations differed significantly for average depth, floral color was not fixed within most populations; in fact, 85% of populations exhibited at least three of the four color classes. Mean flower area ranged from 2.06 cm^2^ to 3.59 cm^2^ across populations (F_19_ = 10.08, P < 0.001; Figure 4, Tables S5– S6)–a difference of ∼70% between smallest and largest flowered populations–and average values of petal width ranged from 0.69 cm to 1.03 cm across populations (F_19_ = 11.43, P < 0.001; Figure 4, Tables S5–S6). While the proportional variation in petal width was less than that of flower area, the differences are still striking, with widest-petaled populations having petals ∼50% broader than the narrowest, again on average.

**Figure 4.**
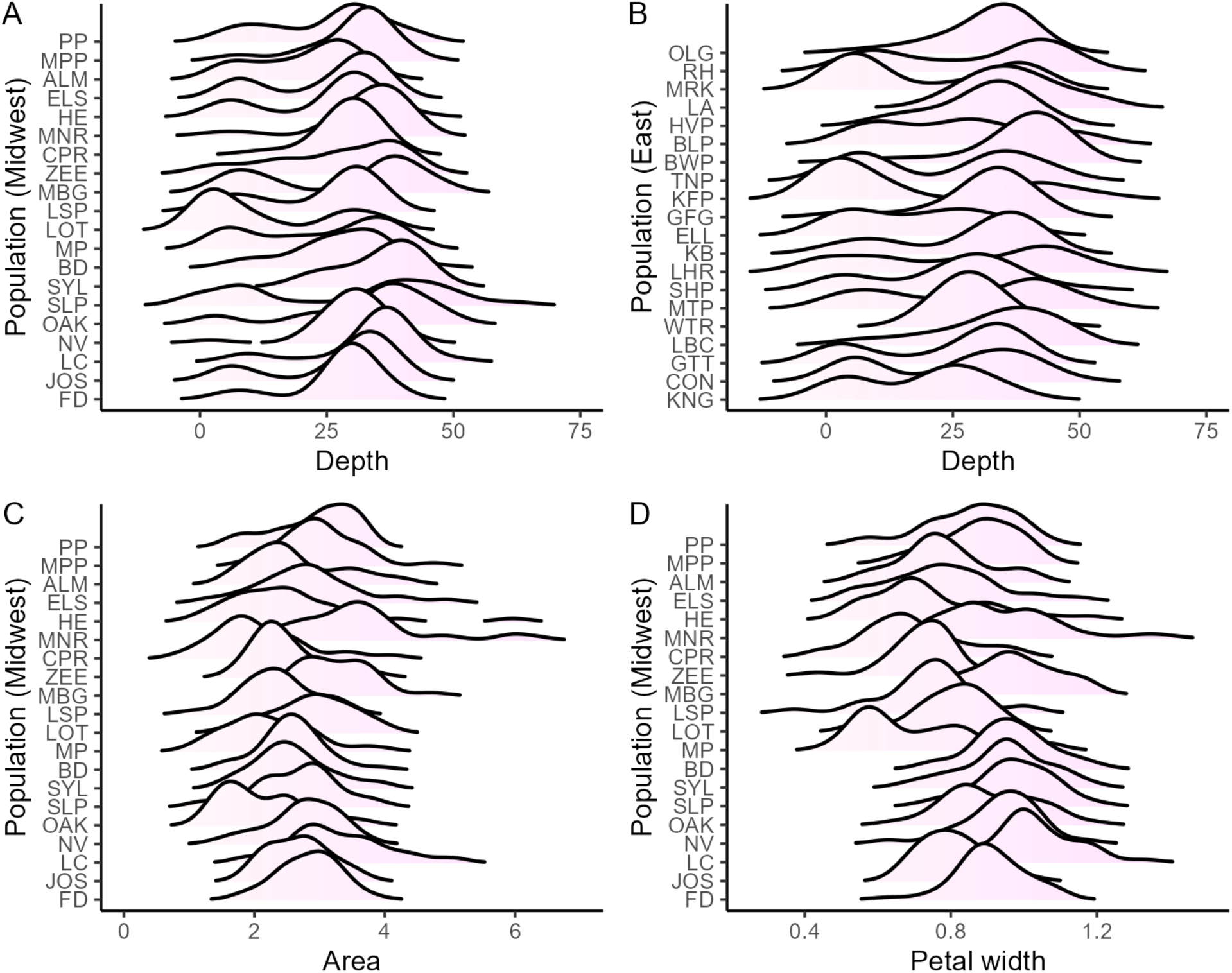
Frequency histograms of the three floral traits (color, size, shape) highlight apparent variation both within and across populations. A) Floral color (Depth) in the Midwest and B) East transects. Smaller depth values capture white and pale floral colors whereas darker values capture pink or purple colors. C) Floral size (Area, cm^2^) and D) petal width (cm) measured in populations from the Midwest transect. Populations in each panel are ordered by latitude. Average values of each trait for each population are presented in Table S5.

Taken together, these results demonstrate that *H. matronalis* exhibits substantial variation in floral color, floral size, and petal shape both among individuals and among populations. The divergence in floral color (depth) is especially pronounced, with average values of the darkest-flowered populations nearly twice that of the lightest. Variation in flower area is also striking, with extreme populations lying >2 SD from the species mean, while petal width, though more modest compared to floral depth and area, still shows pronounced among-population differentiation.

### Neutral genetic structure

We found low genetic differentiation among populations (F_ST_ = 0.016, 95% CI: 0.013-0.018) and little evidence for neutral genetic structure between transects (Midwest vs East F_ST_ = 0.004, 95% CI: 0.003-0.005). An AMOVA indicated that most genetic variation (93.4%) was within populations, with only 3.3% partitioned among transects or among populations within transects. Although low in magnitude, differentiation among populations and transects was greater than expected under panmixia (both *P* = 0.001). Similar results were obtained when SNP calls were analyzed separately by transect (data not shown).

Admixture analysis similarly supported a single genetic cluster (K = 1) in both nonspatial and spatial models (Table S7). The nonspatial model provided a better fit, with mean cross-validation error closer to zero (–0.43) than the spatial model (–0.59). Examination of K = 2 also revealed no clear pattern of admixture by transect or latitude (Figure S4).

Despite the lack of strong population structure, genetic isolation by distance was evident when all populations were considered together (r = 0.358, p < 0.001), but not within individual transects (Midwest: r = 0.096, p = 0.563; East: r = 0.179, p = 0.164). Observed heterozygosity was not correlated with latitude (r = 0.02, p = 0.918), but was positively associated with longitude (r = 0.63, p < 0.001), reflecting higher heterozygosity in the East than the Midwest. Mean H_O_ (±SE) was 0.317 ± 0.005 in the Midwest and 0.348 ± 0.003 in the East (Student’s t = 3.88, df = 33, *P* < 0.001).

### P_ST_-F_ST_ comparisons

We compared phenotypic population structure (P_ST_) to neutral genetic differentiation (F_ST_) to assess whether divergence in traits among populations was more consistent with selection or drift. For all floral traits—color (Depth), size (Area), and petal width—(P_ST_) exceeded F_ST_ under most scenarios, indicating that divergence in floral phenotype is more likely driven by selection than by drift (Figure 5). Only in the most conservative case (Depth at *c/h*^2^ = 0.2) did the 95% confidence intervals of (P_ST_) overlap with F_ST_.

**Figure 5.**
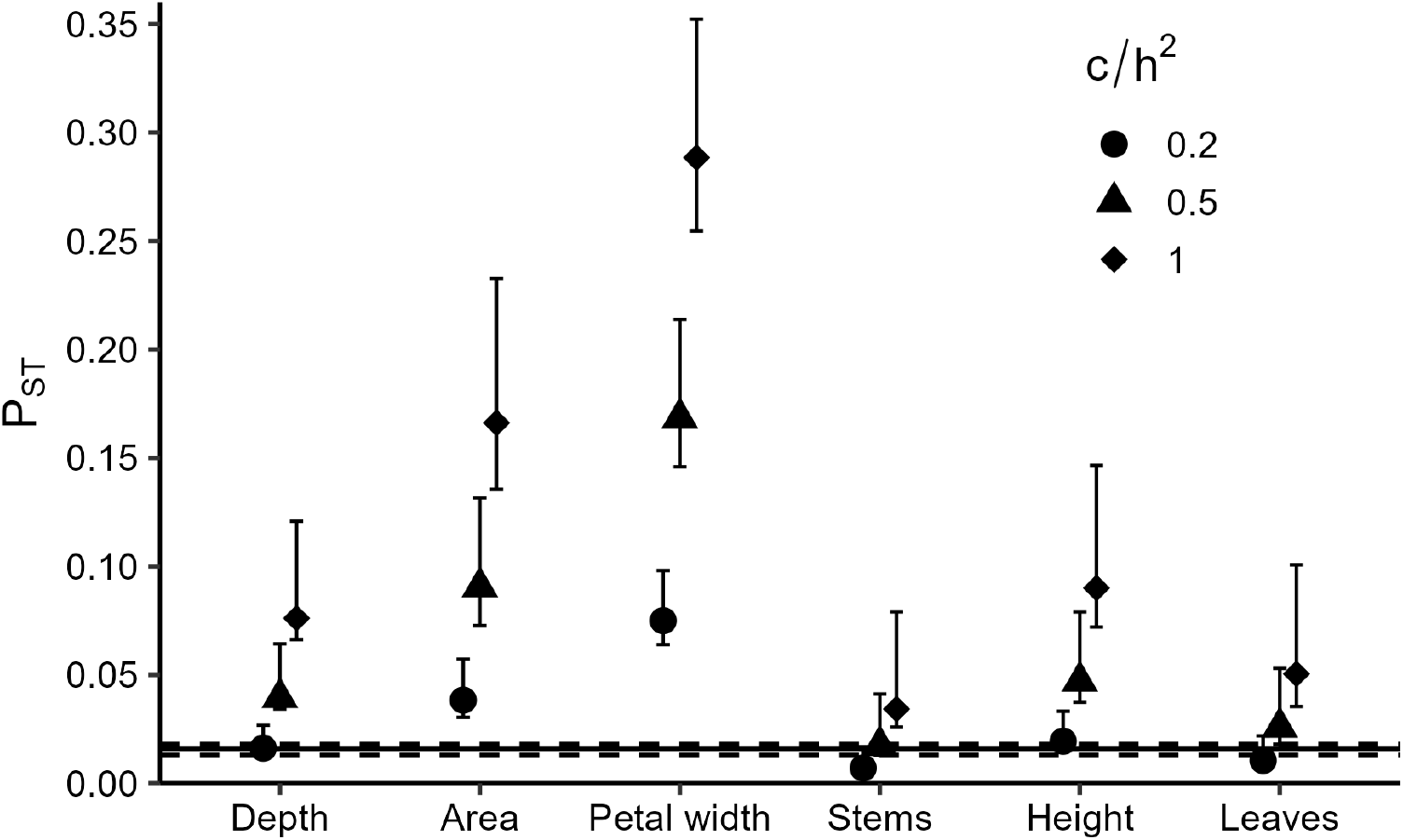
A comparison of the phenotypic structure (P_ST_) of floral color (Depth), floral size (Area), and petal width to the global value of genetic structure (F_ST_) for *Hesperis matronalis*. The P_ST_ values of three plant size traits (height of the tallest stem, stem number, and leaf number) are presented for comparison. Because the ratio of among-population additive genetic variance (c) to within-population heritability (h^2^) is unknown, P_ST_ values and their confidence intervals are shown for potential c/h^2^ values of 0.2, 0.5, and 1.0, with 1.0 representing the least and 0.2 representing the most conservative value, respectively. Population genetic structure (F_ST_) was calculated from the allele frequencies of 692 SNPs, and is represented as the solid horizontal line in the figure with dashed lines representing 95% confidence intervals.

In contrast, evidence for selection on vegetative size traits (stem number, height, leaf number) was less consistent. When *c/h*^2^ = 1, all three traits showed P_ST_ > F_ST_. At *c/h*^2^ = 0.5, plant height and leaf number still exhibited P_ST_ > F_ST_, but in all other comparisons P_ST_ was similar to F_ST_.

Pairwise population comparisons reinforced these patterns. We found evidence for diversifying selection (P_ST_ > F_ST_, non-overlapping CIs) in 34/120 population pairs for flower area, 61/120 for petal width, and 107/630 for depth (which had more comparisons due to being measured across both transects).

### Clinal variation and geographically variable selection

We found no indication of a geographic cline in any trait, but evidence that some floral traits varied with the environment. More specifically, none of the traits exhibited isolation by distance (*i.e.*, nonsignificant Mantel tests, Table S8) indicating there are no visible geographic clines for the examined traits. We found a moderate positive correlation (r = 0.35, *P* = 0.029) between floral color (depth) and the average June minimum temperature (June_avg_tmin; Table S8), but this relationship was no longer significant following corrections for multiple tests (*P* = 0.144). Petal width significantly varied according to the average June minimum temperature (r = 0.61, *P* = 0.02; Table S8), with wider petals (*e.g.,* top row, Figure 2) present in sites that exhibited higher values of average June minimum temperature. This association remained after multiple test corrections.

Multiple regression of distance matrices (MRM), which accounts for both geographic distance and neutral genetic differentiation, likewise revealed a positive association between floral color (depth) and June minimum temperature (June_avg_tmin; β ± SE = 0.14 ± 0.01, P = 0.013; Table S9). As with the Mantel tests, this relationship did not remain significant after correction for multiple comparisons. In contrast, MRM analysis showed that petal width was positively associated with June minimum temperature (β = 0.34 ± 0.007, P = 0.035; Table S9), and again this relationship remained significant following correction. This pattern suggests that aspects of floral morphology, particularly petal width, are shaped by environmental selection linked to thermal conditions, which potentially contributes to the maintenance of phenotypic variation across populations despite high within-population diversity.

### Phenotypic selection

We used Lande–Arnold selection analyses to test for selection on floral and vegetative traits in contemporary *Hesperis matronalis* populations. Across all populations, there was no evidence for directional selection on floral traits, but strong positive selection on plant height (β = 0.30 ± 0.04, *P* < 0.001) and stem number (β = 0.44 ± 0.04, *P* < 0.001; Table S10). Although direct selection on color, flower size, or petal width was not detected, we found significant interactions between traits indicating correlative selection. These included interactions between stem number and flower color (γ = 0.49 ± 0.09, *P* < 0.001), height and flower color (γ = –0.27 ± 0.09, *P* = 0.002), and a three-way interaction between stem number, height, and flower color (γ = –0.22 ± 0.05, *P* < 0.001; Table S10). A three-way interaction between petal width, stem number, and flower color was marginally non-significant (γ = –0.26 ± 0.16, *P* = 0.09; Table S10).

Separate analyses for light- (white, pale) and dark-flowered (pink, purple) plants revealed distinct patterns of selection. In dark-flowered plants, there was strong positive selection on both height (β = 0.37 ± 0.04, *P* < 0.001; Table 1) and stem number (β = 0.29 ± 0.04, *P* < 0.001). In contrast, light-flowered plants showed positive selection on stem number (β = 0.81 ± 0.08, *P* < 0.001) but not on height (β = 0.002 ± 0.09, *P* = 0.98). Light-flowered plants also exhibited divergent selection on stem number (γ = 0.16 ± 0.14, *P* < 0.001), with fitness peaks at both average (∼2) and high (∼5) stem numbers, consistent with the multiple optima identified in the fitness surface analyses.

**Table 1.** Linear and non-linear selection acting on floral size (area), petal width and plant size traits (height, stem number) according to floral color groups (dark, light). Linear (directional) selection gradients (β) were determined from models containing only the linear terms, whereas we estimated nonlinear selection gradients (γ) by doubling quadratic regression coefficients in a full model that contained linear terms, quadratic terms, and the cross-product terms of focal traits.

| Trait | Dark |  | Light |  |
| --- | --- | --- | --- | --- |
| | $\beta \pm (se)$ | P | $\beta \pm (se)$ | P |
| Height | **0.371 (0.037)** | **<0.001** | 0.002 (0.095) | 0.98 |
| Stem number | **0.289 (0.039)** | **<0.001** | **0.811 (0.083)** | **<0.001** |
| Area | -0.026 (0.056) | 0.64 | 0.125 (0.145) | 0.39 |
| Petal width | 0.097 (0.056) | 0.59 | -0.198 (0.148) | 0.19 |
| | $\gamma \pm (se)$ | P | $\gamma \pm (se)$ | P |
| Height x Height | 0.006 (0.071) | 0.93 | -0.245 (0.271) | 0.29 |
| Stem number x Stem number | -0.099 (0.068) | 0.146 | **0.161 (0.149)** | **<0.001** |
| Area x Area | 0.067 (0.063) | 0.286 | 0.422 (0.080) | 0.27 |
| Petal width x Petal width | -0.097 (0.066) | 0.146 | -0.258 (0.232) | 0.29 |
| Height x Stem number | 0.007 (0.039) | 0.861 | **-0.279 (0.065)** | **<0.001** |
| Stem number x Petal width | 0.061 (0.044) | 0.171 | **-0.2 (0.093)** | **0.03** |
| Height x Petal width | 0.041 (0.040) | 0.307 | 0.137 (0.096) | 0.16 |

Patterns of correlative selection also differed between color types. For light-flowered plants, we uncovered negative correlational selection between height and stem number (γ = –0.28 ± 0.07, *P* < 0.001; Figure 6), with highest fitness in shorter (∼53 cm), many-stemmed (>4) individuals. Additionally, light-flowered plants showed negative correlational selection between petal width and stem number (γ = – 0.20 ± 0.09, *P* = 0.03; Figure 6), with highest fitness in plants having many stems and petal widths slightly below the mean (∼0.80 cm vs. 0.88 cm). Dark-flowered plants showed no evidence for correlational selection among measured traits.

**Figure 6.**
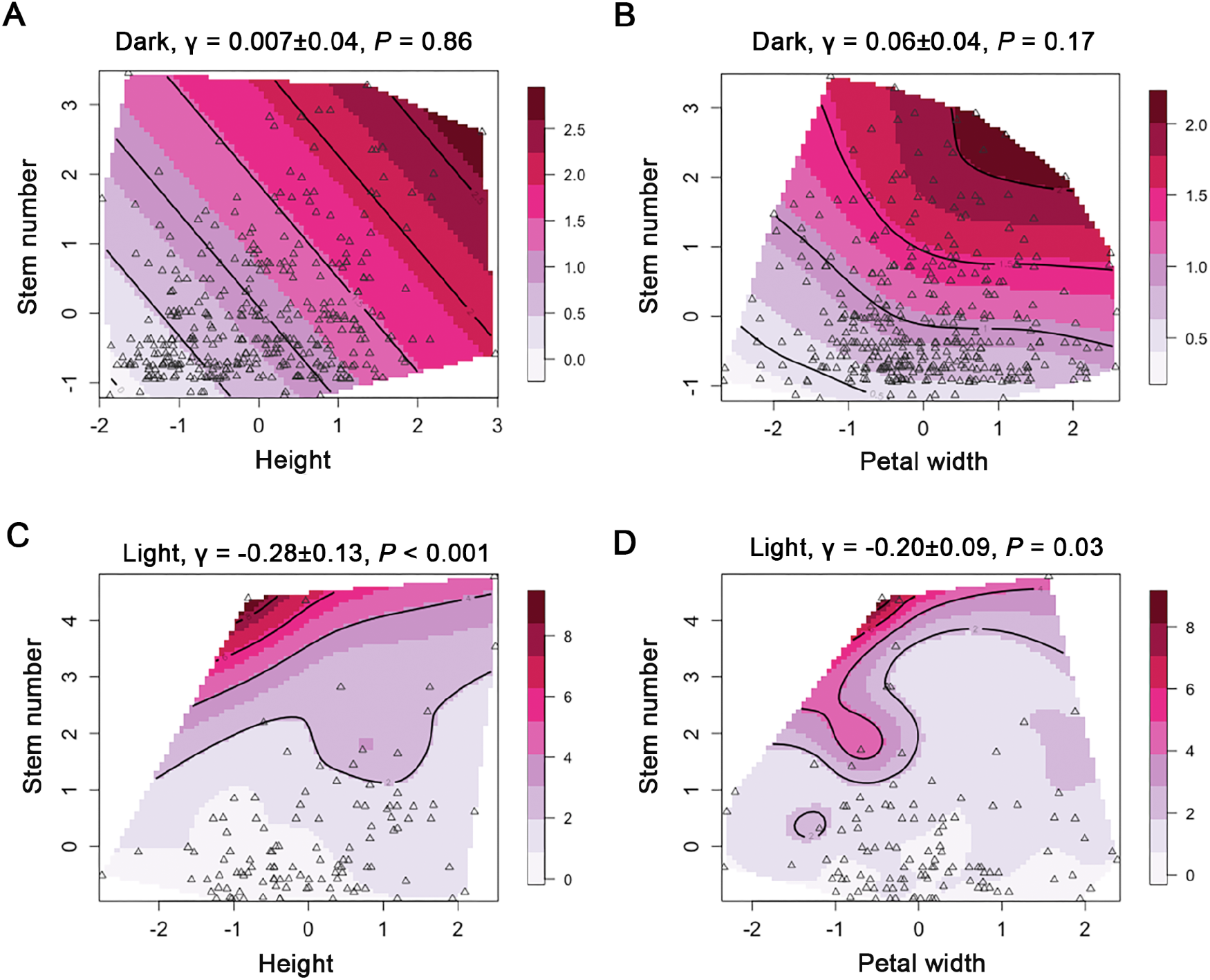
Different peaks of high fitness for dark and light floral phenotypes indicate that different combinations of traits are favored according to floral color. Darker flowered individuals (pink, purple) exhibited evidence for positive selection on both stem number and height, but did not show evidence of correlative selection between these traits (A) or between stem number and petal width (B) whereas plants with light flowers (white, pale) exhibited negative correlational selection between height and stem number (C), and correlational selection between stem number and petal width (D).

Despite these differences in selection gradients, we found no evidence that mean trait values differed between flower color classes. Plant height, stem number, leaf number, flower area, and petal width did not differ significantly between light- and dark-flowered plants (Figure S5, Table S11).

## Discussion

Our comprehensive approach assessing floral variation in *Hesperis matronalis* suggests that floral traits are under the influence of selection, but that the separate traits are likely responding to different selective forces. Perhaps most intriguing is the evidence that populations exhibit significant trait differentiation despite little genetic structure, with phenotypic differentiation exceeding that based on neutral genetic variation. Below, we discuss patterns of trait variation across the landscape, the evidence supporting selection, and potential indications of within-population dynamics that may underlie divergence in floral trait values between populations.

### *Extensive floral variation within* Hesperis matronalis

*Hesperis matronalis*, an introduced weed present in the US since ∼1840, is best known for its striking floral color variation. Our survey of 40 populations confirms prior observations that floral color is variable within most populations: only one population was monomorphic (WTR, STable 1) while the vast majority contained at least three perceptible color classes. Analyses indicate that floral color in this species exhibits multiple distinct peaks with some overlap among trait values, a pattern similar to that seen in *Claytonia virginica* (Frey 2004, 2007) and the common morning glory, *Ipomoea purpurea* (Rausher and Fry 1993; Clegg and Durbin 2000). Like *H. matronalis, I. purpurea* produces blue (purple), pink, or white flowers with varying intensities, which are controlled by at least four loci: one major locus determining pink or blue (purple) flowers and additional loci modifying intensity or producing white flowers epistatically (Clegg and Durbin 2000). Unlike *I. purpurea*, however, *H. matronalis* flowers do not exhibit strong variation in intensity or pigmented floral tubes, and while the genetic basis of *H. matronalis*’s major floral color classes remains unresolved, evidence suggests that two loci may underlie the purple–white polymorphism (Majetic 2008b). Further work will be necessary to determine the genetic control underlying the other major color classes as well as the influence of the environment on the intermediate phenotypes bridging the four major classes.

Our study also revealed striking diversity in floral size and petal width within *Hesperis matronalis* populations. The largest flowers were seven times larger than the smallest, and petal widths varied 3.7-fold across individuals. For both traits, among-individual variation exceeded within-individual variation, consistent with plants producing flowers of uniform size. Within-population variation in floral size traits is widely reported in other species (Galen 1999b; Worley et al. 2000) and often linked to pollinator preferences (Galen and Cuba 2001; Elle and Carney 2003; Strauss and Whittall 2006), though explicit documentation of petal size variation is less common (Parachnowitsch and Kessler 2010; Koski 2023). Further, work in other species, such as the related *Brassica rapa*, shows that variation in floral size has a heritable genetic basis and responds readily to artificial selection (Sarkissian and Harder 2001; Gervasi and Schiestl 2017), with overall floral size and petal width even responding indirectly to selection on floral volatiles (Zu et al. 2016). While our study was not designed to quantify heritability in floral size and petal width in *H. matronalis*, the substantial variation in floral size traits that we have documented likely has a heritable component that is responsive to selection. Future studies designed to disentangle the influence of the environment from the genetics underlying floral size traits in this species are needed.

Just as striking as the high within-population variation in floral traits, *Hesperis matronalis* populations exhibited wide variation among populations in mean trait values: for example, the average floral color depth of the darkest population was 190% greater than that of the lightest population, and floral shape and petal width likewise showed pronounced among-population variation. This among-population differentiation in trait values was not associated with spatial distance, such that there are no perceptible geographic clines in any floral trait examined. Unlike many plant species, this species shows no evidence of isolation by distance in floral color, a striking departure from the typical geographic structuring of floral traits. For example, clinal variation in floral color is reported for *Silene vulgaris* (Berardi et al. 2016), for floral reflectance plasticity in *Plantago lanceolata* (Marshall et al. 2019), and for the pink floral color locus in *Ipomoea purpurea* (Epperson and Clegg 1986). More abrupt clines in floral color are seen in *Mimulus aurantiacus* (Streisfeld and Kohn 2005) and *Linanthus parryae* (Schemske and Bierzychudek 2007), which exhibit sharp differentiation of red versus yellow flowers between inland and coastal regions of California, and an exceptionally steep transition from white to blue flowers within a 30-m ravine, respectively.

### *Floral variation covaries with environment in* Hesperis matronalis

While we found no evidence of a geographic cline in traits, we found some indication, albeit weak, that floral color variation in *Hesperis matronalis* may be shaped by an environmental cline. Specifically, we found a moderate positive relationship between minimum June nighttime temperature and floral depth, suggesting that populations flowering under warmer nocturnal conditions exhibit darker morphs. While darker flowers are associated with cooler environments or higher elevations in some species (e.g., *Campanula americana*, where warmer sites had lighter petals), work in *Ipomoea purpurea* demonstrates that darker morphs can persist or even perform well under warmer conditions (Coberly and Rausher 2003a). Thus, relationships between floral color and environment are likely to be species-specific. We note that the observed association between floral color and temperature in *H. matronalis* was not significant after multiple test corrections, such that additional studies will be required to determine whether environmental factors truly shape floral color variation in this species.

In contrast, we uncovered a strong relationship between petal width and the average minimum June nighttime temperature, a relationship that remained significant following corrections for multiple tests. Specifically, populations experiencing warmer June nights tend to produce wider petals, suggesting that warmer night time temperatures induce larger petal widths, or that wider petals may be adaptive in areas with higher nighttime temperatures. We did not detect any environmental association with overall floral area, despite a strong correlation between petal width and total floral area (r = 0.7). Notably, both large and small flowers in this species can possess either wide or narrow petals, indicating that overall floral size is not strictly determined by petal width variation. Our findings contrast somewhat to other species where overall flower size varies along altitudinal and/or climatic gradients. For example, *Campanula rotundifolia* populations at higher altitudes produce larger corollas (Maad et al. 2013), and in *Plantanthera dilatata*, floral size traits correlate with temperature gradients (Plendl et al. 2024). Thus, thermal environment may be an important driver of floral size variation in plants, but the specific trait responding—whether overall floral size or a component trait such as petal width—is likely species dependent.

### Patterns of among-population variation in floral traits support selection

Our examination of population genetic structure found low genetic differentiation among populations and only slight differentiation between transects. We further found no evidence of isolation by distance within either transect, suggesting a near absence of spatial genetic differentiation across the landscape. In contrast, phenotypic differentiation of floral traits exceeded that of neutral genetic differentiation (i.e., P_ST_ > F_ST_) under most scenarios, indicating that trait variation is influenced by selection. To approximate Q_ST_, we compared P_ST_ and F_ST_ under three scenarios of c/h^2^ (0.2, 0.5, and 1.0), where c is the proportion of phenotypic variance due to additive effects among populations and h^2^ is the additive genetic variance within populations. At the null expectation of 1 (c = h^2^), all traits examined showed P_ST_ > F_ST_, consistent with divergent selection. At the most conservative scenario (c/h^2^ = 0.2), only petal width and overall floral size P_ST_ exceeded F_ST_, suggesting that most other traits are influenced by drift. Floral color (depth) showed P_ST_ > F_ST_ at intermediate and null scenarios (0.5 and 1), and height followed a similar pattern, whereas the other vegetative traits generally did not exceed F_ST_.

Overall, our results indicate that while most vegetative traits show limited evidence for selection, floral size, petal width, floral color, and height exhibit signals of divergent selection across the landscape. Although the number of studies directly comparing floral phenotypic and neutral genetic divergence remains limited, our findings are consistent with patterns reported in other plant systems. For example, both floral color and floral size show evidence of adaptive divergence in *Ipomopsis aggregata* (Milano et al. 2016), while divergent selection has also been detected for floral reflectance in *Plantago lanceolata* (Marshall et al. 2019) and floral color in *Silene vulgaris* (Berardi et al. 2016). Despite the similarity across systems, we note that P_ST_–F_ST_ comparisons like the one we present here must be interpreted cautiously, and that direct estimation of Q_ST_ will require heritability estimates from populations grown in a common garden—a challenging but critical step for biennial species such as *H. matronalis*.

### Patterns of contemporary selection differ by floral morphologies

Patterns of phenotypic selection in contemporary populations of *Hesperis matronalis* show that vegetative traits are under directional selection, with strong fitness advantages for taller, many-stemmed individuals. Although we did not find evidence of direct selection on any floral trait, floral color and petal width showed strong correlative and context-dependent selection, particularly in light-flowered plants where disruptive and correlational selection shape fitness peaks at multiple trait combinations. More specifically, light-flowered individuals exhibited a fitness peak when shorter and many-stemmed, or many-stemmed with petal widths slightly below the mean. These findings highlight a complex selective landscape in which vegetative performance traits influence overall fitness, while floral color interacts with plant architecture and petal size traits to mediate the form of selection.

Evidence for linear selection on floral color, size, and shape has been reported in many plant species summarized in (Caruso et al. 2019), typically in the context of pollinator- or antagonist-mediated selection. Further, pollinators often favor particular combinations of traits—for example, taller *Erysimum mediohispanicum* plants with more flowers and longer petals (Gómez 2003), or shorter floral displays with smaller petals in *Trillium discolor* (Koski 2023) are preferred. Whether such trait combinations lead to nonlinear or correlational selection is less well understood, as quadratic selection gradients are reported far less frequently than linear gradients on individual traits (Caruso et al. 2019; Koski 2023).

Nevertheless, available studies demonstrate that fitness peaks can emerge from specific trait combinations, such as increased stalk height with smaller flowers in *E. mediohispanicum* (Gómez 2003) or shorter stalks with smaller flowers in *T. discolor (Koski 2023)*. Disruptive and correlative selection would therefore be expected to generate variable floral morphs within populations, and could potentially reflect the differing preferences of distinct members of the pollination community. While we have not yet performed pollinator observations on *Hesperis matronalis*, the presence of fitness peaks for combinations of traits could indicate that a variable pollinator community, with different preferences, are acting to maintain variation in floral traits within populations (Brothers and Atwell 2014; Klecka et al. 2018).

## Conclusions

Our findings demonstrate that *Hesperis matronalis* harbors extensive variation in floral traits both within and among populations, with divergence in color, size, and petal width exceeding neutral genetic differentiation and thus likely reflecting the action of selection. While vegetative traits appear to be the main targets of directional selection, floral color and size traits show signatures of divergent, disruptive, and correlative selection, particularly in ways that may reflect the heterogeneous preferences of pollinators. The absence of general patterns of genetic isolation by distance, coupled with strong trait differentiation, suggests that selection rather than drift underlies much of the observed variation. Together, these results highlight a complex landscape in which multiple evolutionary forces act to maintain floral diversity in this species.

An important caveat to our conclusions is that we do not yet know how responsive these floral traits are to environmental conditions. While our field surveys suggest a genetic basis to the observed variation, greenhouse experiments will be essential to disentangle genetic effects from environmental influences and to more precisely determine the extent to which floral color and morphology are plastic across environments. Such work is particularly important for biennial species like *Hesperis matronalis,* where environmental variation across years may mask or amplify genetic differences. Nonetheless, our study provides evidence of extensive trait variation, population differentiation exceeding neutral expectations, and signatures of selection shaping floral and vegetative traits. Future work should focus on estimating heritability in a common garden framework and linking trait variation to pollinator behavior, steps that are critical for fully resolving the evolutionary processes maintaining floral diversity in this species.

## Tables

### Figures

Supplementary information for: Color, size, shape: The drivers of floral variation in *Hesperis matronalis* (Dame’s Rocket)

**TABLE S1.** Population characteristics of *H. matronalis* sites sampled across two transects in 2023. The geographic location and the number of phenotyped and genotyped individuals for each site are provided as well as observed and expected heterozygosity of each population.

| Transect | Site | Longitude | Latitude | Elevation (m) | N phenotyped |  |  | N genotyped | Heterozygosity |  |
| --- | --- | --- | --- | --- | --- | --- | --- | --- | --- | --- |
|  |  |  |  |  | Vegetative traits | Flower color | Flower shape |  | Ho | He |
| Midwest | FD | -82.115 | 39.703 | 237 | 48 | 30 | 48 | 0 | -- | -- |
|  | JOS | -82.393 | 39.874 | 319 | 36 | 30 | 36 | 2 | 0.272 | 0.204 |
|  | LC | -82.609 | 40.268 | 333 | 42 | 30 | 41 | 10 | 0.319 | 0.144 |
|  | NV | -82.733 | 40.783 | 352 | 30 | 30 | 30 | 10 | 0.308 | 0.140 |
|  | OAK | -83.686 | 41.022 | 237 | 36 | 30 | 36 | 10 | 0.288 | 0.134 |
|  | SLP | -83.66 | 41.362 | 212 | 33 | 28 | 30 | 10 | 0.315 | 0.142 |
|  | SYL | -83.702 | 41.713 | 209 | 33 | 30 | 32 | 10 | 0.334 | 0.153 |
|  | BD | -83.927 | 41.993 | 241 | 36 | 30 | 36 | 10 | 0.332 | 0.151 |
|  | MP | -83.771 | 42.287 | 286 | 30 | 30 | 30 | 10 | 0.335 | 0.154 |
|  | LOT | -83.722 | 42.288 | 248 | 30 | 30 | 30 | 9 | 0.317 | 0.147 |
|  | LSP | -83.743 | 42.292 | 248 | 30 | 30 | 30 | 8 | 0.295 | 0.136 |
|  | MBG | -83.666 | 42.295 | 252 | 36 | 30 | 36 | 10 | 0.337 | 0.151 |
|  | ZEE | -83.841 | 42.325 | 264 | 35 | 30 | 35 | 10 | 0.346 | 0.156 |
|  | CPR | -83.756 | 42.534 | 295 | 30 | 30 | 30 | 0 | -- | -- |
|  | MNR | -84.416 | 42.747 | 279 | 40 | 30 | 40 | 10 | 0.325 | 0.146 |
|  | HE | -83.617 | 42.794 | 279 | 35 | 30 | 35 | 3 | 0.292 | 0.203 |
|  | ELS | -84.382 | 43.093 | 218 | 30 | 30 | 30 | 9 | 0.288 | 0.133 |
|  | ALM | -84.656 | 43.377 | 222 | 40 | 30 | 40 | 10 | 0.291 | 0.136 |
|  | MPP | -84.787 | 43.598 | 223 | 35 | 30 | 35 | 2 | 0.323 | 0.136 |
|  | PP | -84.768 | 43.813 | 264 | 35 | 30 | 35 | 10 | 0.347 | 0.158 |
| East | KNG | -79.269 | 39.899 | 590 | - | 30 | - | 5 | 0.383 | 0.168 |
|  | CON | -78.86 | 40.27 | 608 | - | 30 | - | 9 | 0.352 | 0.160 |
|  | GTT | -78.731 | 40.482 | 618 | - | 30 | - | 8 | 0.352 | 0.160 |
|  | LBC | -78.679 | 40.835 | 401 | - | 30 | - | 4 | 0.355 | 0.157 |
|  | WTR | -78.787 | 41.173 | 465 | - | 30 | - | 5 | 0.351 | 0.157 |
|  | MTP | -78.725 | 41.422 | 434 | - | 30 | - | 10 | 0.339 | 0.156 |
|  | SHP | -78.649 | 41.906 | 457 | - | 30 | - | 9 | 0.356 | 0.163 |
|  | LHR | -78.607 | 41.957 | 466 | - | 30 | - | 6 | 0.341 | 0.155 |
|  | KB | -78.679 | 42.153 | 423 | - | 30 | - | 10 | 0.352 | 0.160 |
|  | ELL | -78.67 | 42.274 | 467 | - | 30 | - | 7 | 0.336 | 0.154 |
|  | GFG | -78.683 | 42.504 | 411 | - | 30 | - | 10 | 0.360 | 0.163 |
|  | KFP | -78.637 | 42.768 | 270 | - | 30 | - | 5 | 0.325 | 0.146 |
|  | TNP | -78.854 | 42.846 | 175 | - | 30 | - | 9 | 0.360 | 0.162 |
|  | BWP | -78.739 | 43.007 | 176 | - | 30 | - | 10 | 0.346 | 0.157 |
|  | BLP | -78.929 | 43.184 | 175 | - | 30 | - | 5 | 0.351 | 0.157 |
|  | HVP | -79.865 | 43.303 | 82 | - | 30 | - | 9 | 0.352 | 0.161 |
|  | LA | -79.851 | 43.526 | 198 | - | 30 | - | 5 | 0.347 | 0.157 |
|  | MRK | -79.403 | 43.836 | 200 | - | 30 | - | 5 | 0.352 | 0.157 |
|  | RH | -79.458 | 43.879 | 187 | - | 30 | - | 10 | 0.315 | 0.148 |
|  | OLG | -79.541 | 43.951 | 302 | - | 30 | - | 5 | 0.341 | 0.152 |

**FIGURE S1.**
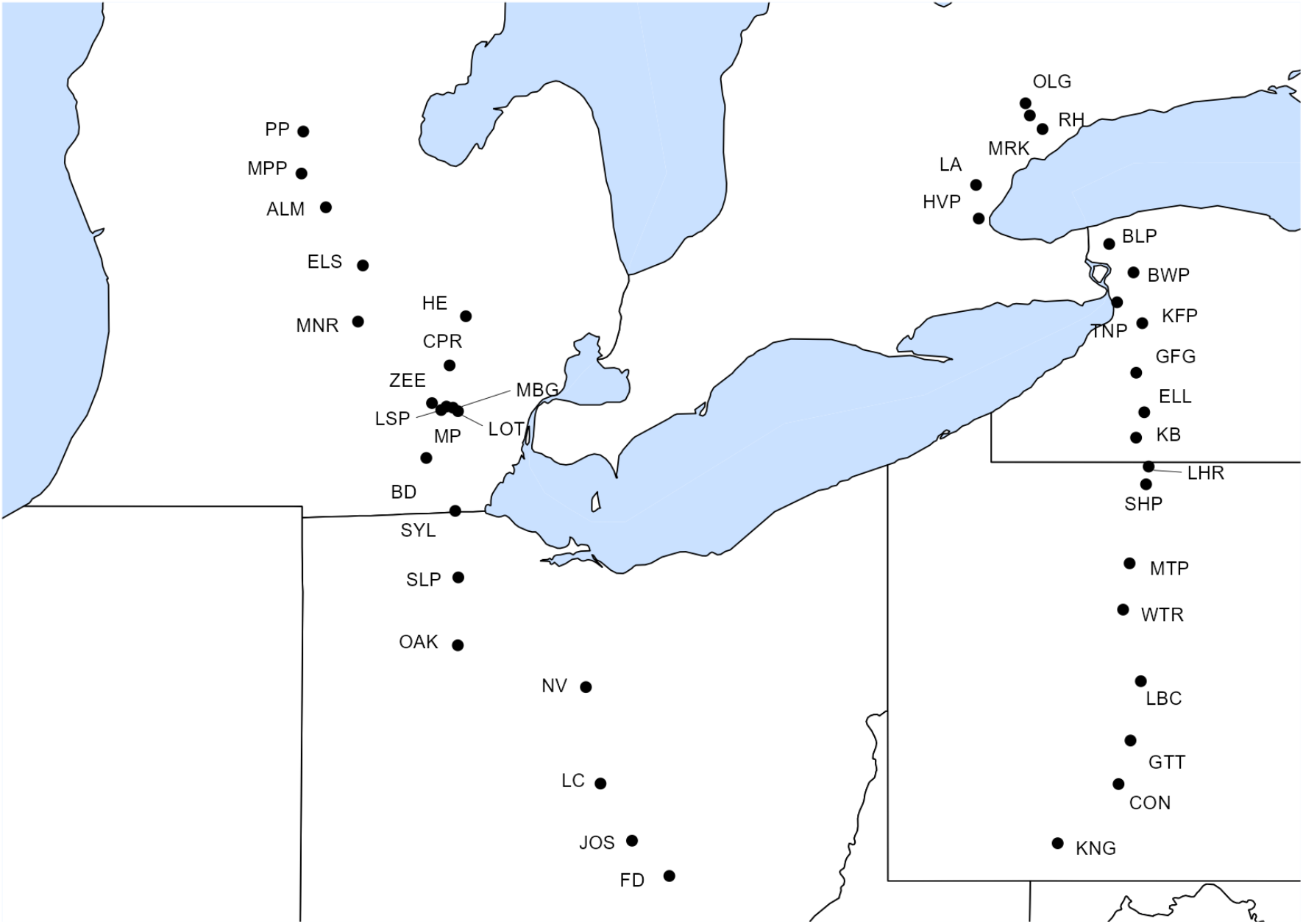
Map of sites sampled in 2023. Forty sites were sampled in total, with 20 from each of two transects. Descriptive information for each location is provided in Table S1.

## Supplemental methods 1

### Environmental variables analysis

We performed PCA to condense the 19 WorldClim bioclimatic variables into a few axes of environmental variation in floral traits across our sites. We first performed this analysis using all population locations across both transects to examine potential environmental correlates associated with flower color (depth). Among all midwest and eastern sites visited, the first two principal components (PCs) of the 19 Bioclim variables collectively explained 78% of the variance (Figure S2). Factor loadings indicated that PC1 loads negatively on temperature seasonality (bio4), max temperature of the warmest month (bio5), and mean temperature of the warmest quarter (bio10), and positively on annual precipitation (bio12), precipitation of driest month (bio14), and coldest quarter precipitation (bio19) (Table S2). PC1 therefore captures a gradient from hotter, more seasonally variable climates (low PC1 values) to cooler, wetter, and more stable climates (high PC1 values). PC2 was primarily associated with cold-season temperatures, exhibiting strong positive loadings on min temperature of coldest month (bio6), mean temperature of coldest quarter (bio11), and temp of driest quarter (bio9) (Table S2). Thus, PC2 reflects how cold the coldest parts of the year are—higher PC2 values mean warmer winters, while lower values mean colder winters.

We next performed PCA analysis using only the midwest sites to examine environmental correlates associated with flower size, shape, and the plant size traits. In this analysis, the first two PCs of the 19 Bioclim variables collectively explained 89% of the variance (Figure S2). Similar to the full dataset, the Midwest PC1 exhibits positive loadings on overall temperatures (*e.g.*, annual mean temp, bio1; min temp of coldest month, bio6), and negative loadings on precipitation seasonality and max temperature of warmest month (Table S2), meaning that Midwest PC1 captures a temperature gradient, with higher values meaning warmer, more stable climates, and lower values meaning cooler, more variable climates. The Midwest PC2 reflects seasonal differences in rainfall, with higher values suggesting warmer, wetter seasons, and lower values reflecting cooler, drier conditions during key parts of the year (*i.e.*, positive loadings on bio8 (mean temp of wettest quarter), bio9 (temp of driest quarter), and negative loadings on precipitation of warmest and wettest quarters).

**FIGURE S2.**
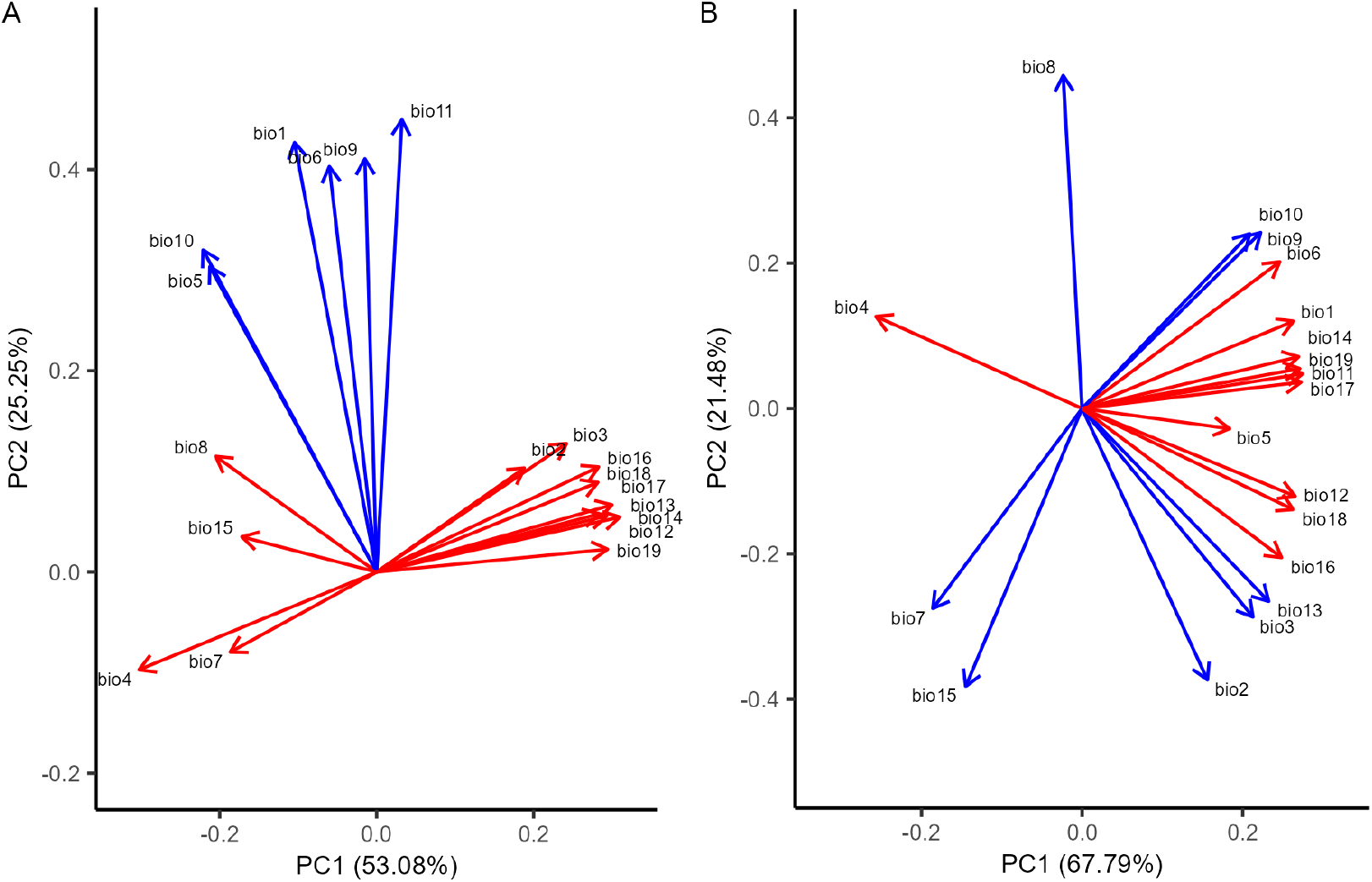
Principal component analysis (PCA) of the 19 WorldClim Bioclimatic variables. Each Bioclimatic variable loading was colored red if it contributed more to the first PCA axis or blue if it contributed more to the second PCA axis. A) PCA for all 40 sites. The first two PCA axes explain 53.08% and 25.25% of the total variance, respectively. WorldClim PC1 was associated with greater precipitation, lower temperature seasonality, and cooler summer high temperatures, and WorldClim PC2 was associated with warmer annual mean temperature and seasonal temperature highs. Results were similar when excluding four sites omitted from F_ST_ analysis due to sample size. B) PCA for our 20 Midwest-transect sites. The first two PCA axes explain 67.79% and 21.48% of the total variance, respectively. WorldClim PC1 was associated with higher annual temperatures, greater precipitation, and reduced temperature seasonality, and WorldClim PC2 was associated with reduced temperature seasonality and daily range as well as precipitation seasonality and precipitation of the wettest three months. When excluding four sites omitted from F_ST_ analysis due to sample size results were similar, except for a reversal in the direction of PC1 loadings.

**TABLE S2.** The results of a principal component analysis used to summarize variation of the WorldClim 19 bioclim variables to composite variables for testing climate. The Eigenvalues of each of the bioclim variables are presented for the two axes that explain the majority of the variation.

|  |  |  | Both Transects |  | Midwest transect |  |
| --- | --- | --- | --- | --- | --- | --- |
|  | Variable | Description | PC1 | PC2 | PC1 | PC2 |
| Proportion Explained |  |  | 0.559 | 0.224 | 0.678 | 0.215 |
| Eigenvalue |  |  | 10.62 | 4.25 | 12.88 | 4.08 |
| Loadings | bio1 | annual mean temperature | -0.16 | 0.39 | 0.263 | 0.12 |
|  | bio2 | mean monthly diurnal range | 0.193 | -0.02 | 0.156 | -0.373 |
|  | bio3 | isothermality ((bio2 / bio7)*100) | 0.243 | 0.041 | 0.213 | -0.287 |
|  | bio4 | temperature seasonality (standard deviation *100) | -0.293 | -0.111 | -0.257 | 0.127 |
|  | bio5 | max temperature of warmest month | -0.244 | 0.203 | 0.184 | -0.028 |
|  | bio6 | min temperature of coldest month | -0.109 | 0.437 | 0.246 | 0.202 |
|  | bio7 | annual temperature range (bio5 - bio6) | -0.168 | -0.226 | -0.186 | -0.275 |
|  | bio8 | mean temperature of wettest quarter | -0.205 | 0.024 | -0.024 | 0.458 |
|  | bio9 | mean temperature of driest quarter | -0.065 | 0.426 | 0.222 | 0.242 |
|  | bio10 | mean temperature of warmest quarter | -0.252 | 0.265 | 0.209 | 0.24 |
|  | bio11 | mean temperature of coldest quarter | -0.016 | 0.472 | 0.275 | 0.048 |
|  | bio12 | annual precipitation | 0.299 | 0.08 | 0.265 | -0.121 |
|  | bio13 | precipitation of wettest month | 0.278 | 0.021 | 0.232 | -0.265 |
|  | bio14 | precipitation of driest month | 0.28 | 0.129 | 0.27 | 0.071 |
|  | bio15 | precipitation seasonality (coefficient of variation) | -0.16 | -0.11 | -0.145 | -0.383 |
|  | bio16 | precipitation of wettest quarter | 0.274 | 0.077 | 0.248 | -0.205 |
|  | bio17 | precipitation of driest quarter | 0.286 | 0.14 | 0.273 | 0.036 |
|  | bio18 | precipitation of warmest quarter | 0.273 | 0.057 | 0.263 | -0.139 |
|  | bio19 | precipitation of coldest quarter | 0.282 | 0.106 | 0.271 | 0.055 |

TABLE S3. Sample names and library information for ddRAD libraries. The number of raw fastq file PE reads and filtered BAM file PE reads that remained for SNP calling are provided for each sample.

- Table is located at “STable_read_info” tab of google spreadsheet.

**FIGURE S3.**
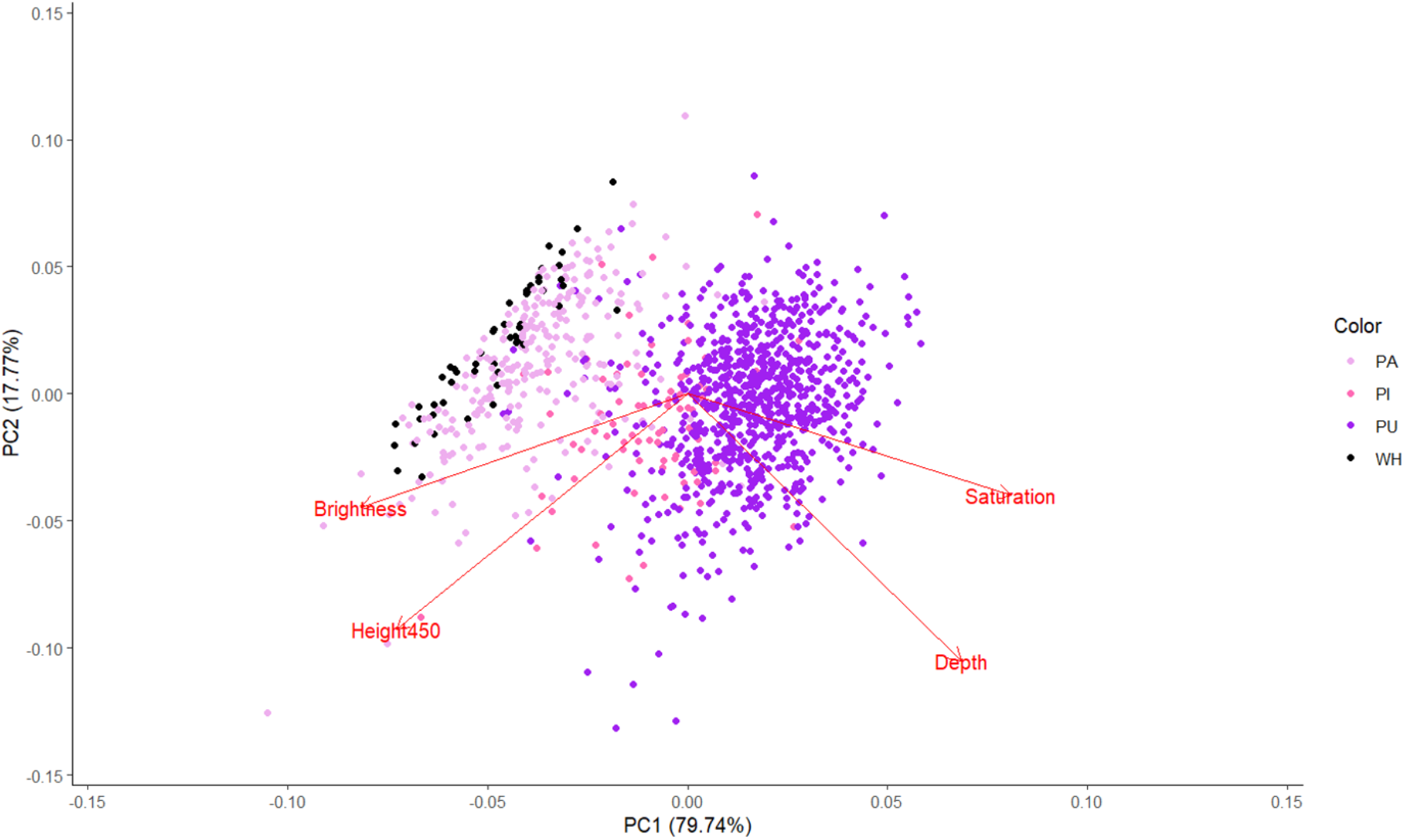
PCA analysis in which we overlaid our estimates of qualitative floral color (WH = white, PU = purple, PI = pink, PA = pale) of each individual with their spectrometry readings (values of depth, saturation, reflectance curve height at 450, brightness).

**TABLE S4.** The results of analyses of variance to determine if flower size (Area) and petal width differed among individuals.

| Traits | Individual ID |  |  |
| --- | --- | --- | --- |
| | Df | $\chi^2$ | P |
| Area | 697 | 10454 | <0.001 |
| Petal width | 697 | 11651 | <0.001 |

**TABLE S5.** Population mean and standard error (se) for flower color (Depth), size (Area), and Petal width. Flower area and petal width were only measured for the midwest transect.

| Site | Depth | Area | Petal width |
| --- | --- | --- | --- |
| FD | 27.68 (1.56) | 2.84 (0.07) | 0.78 (0.12) |
| JOS | 28.02 (2.07) | 2.68 (0.08) | 0.97 (0.10) |
| LC | 33.83 (1.76) | 3.28 (0.11) | 0.69 (0.13) |
| NV | 29.93 (1.29) | 2.76 (0.1) | 0.78 (0.15) |
| OAK | 33.29 (2.32) | 2.09 (0.1) | 0.91 (0.09) |
| SLP | 28.87 (3.67) | 2.67 (0.11) | 0.74 (0.16) |
| SYL | 35.85 (1.38) | 2.57 (0.1) | 0.81 (0.09) |
| BD | 27.57 (1.69) | 2.57 (0.08) | 1.03 (0.11) |
| MP | 23.82 (2.43) | 2.22 (0.12) | 0.80 (0.11) |
| LOT | 13.32 (2.47) | 2.9 (0.11) | 0.74 (0.13) |
| LSP | 23.85 (2.05) | 2.36 (0.1) | 0.97 (0.11) |
| MBG | 28.0 (2.81) | 3.2 (0.09) | 0.94 (0.15) |
| ZEE | 27.16 (2.27) | 2.49 (0.09) | 0.70 (0.16) |
| CPR | 29.19 (1.04) | 2.06 (0.13) | 0.88 (0.10) |
| MNR | 33.06 (1.58) | 3.59 (0.15) | 0.94 (0.10) |
| HE | 25.05 (2.14) | 2.43 (0.15) | 0.91 (0.12) |
| ELS | 25.3 (2.03) | 2.82 (0.14) | 0.85 (0.12) |
| ALM | 20.87 (1.74) | 2.64 (0.11) | 0.98 (0.10) |
| MPP | 30.26 (1.5) | 3.0 (0.09) | 0.95 (0.12) |
| PP | 25.2 (2.08) | 2.93 (0.1) | 0.75 (0.12) |
| KNG | 19.02 (2.19) | - | - |
| CON | 24.86 (2.69) | - | - |
| GTT | 24.07 (2.59) | - | - |
| LBC | 33.6 (2.13) | - | - |
| WTR | 28.75 (1.01) | - | - |
| MTP | 28.61 (3.08) | - | - |
| SHP | 24.51 (2.44) | - | - |
| LHR | 27.07 (3.43) | - | - |
| KB | 28.27 (2.45) | - | - |
| **ELL** | 17.98 (2.28) | - | - |
| **GFG** | 31.06 (1.75) | - | - |
| **KFP** | 16.19 (3.53) | - | - |
| **TNP** | 23.53 (2.87) | - | - |
| **BWP** | 37.83 (1.86) | - | - |
| **BLP** | 26.01 (2.68) | - | - |
| **HVP** | 30.66 (1.63) | - | - |
| **LA** | 37.06 (1.57) | - | - |
| **MRK** | 19.82 (2.94) | - | - |
| **RH** | 29.28 (2.94) | - | - |
| **OLG** | 31.35 (1.6) | - | - |

**TABLE S6.** The result of an analysis of variance to determine if traits vary among populations. Height, stem number, flower area, and petal width were examined only across populations from the Midwest transect whereas depth was examined across transects.

| **Traits** | **Population** |  |  |
| --- | --- | --- | --- |
|  | **Df** | **F-value** | **P** |
| Height | 19 | 6.01 | **<0.001** |
| Stem number | 19 | 2.78 | **<0.001** |
| Flower area | 19 | 11.43 | **<0.001** |
| Petal width | 19 | 19.35 | **<0.001** |
| Flower color (depth) | 39 | 202.05 | **<0.001** |

**TABLE S7.**
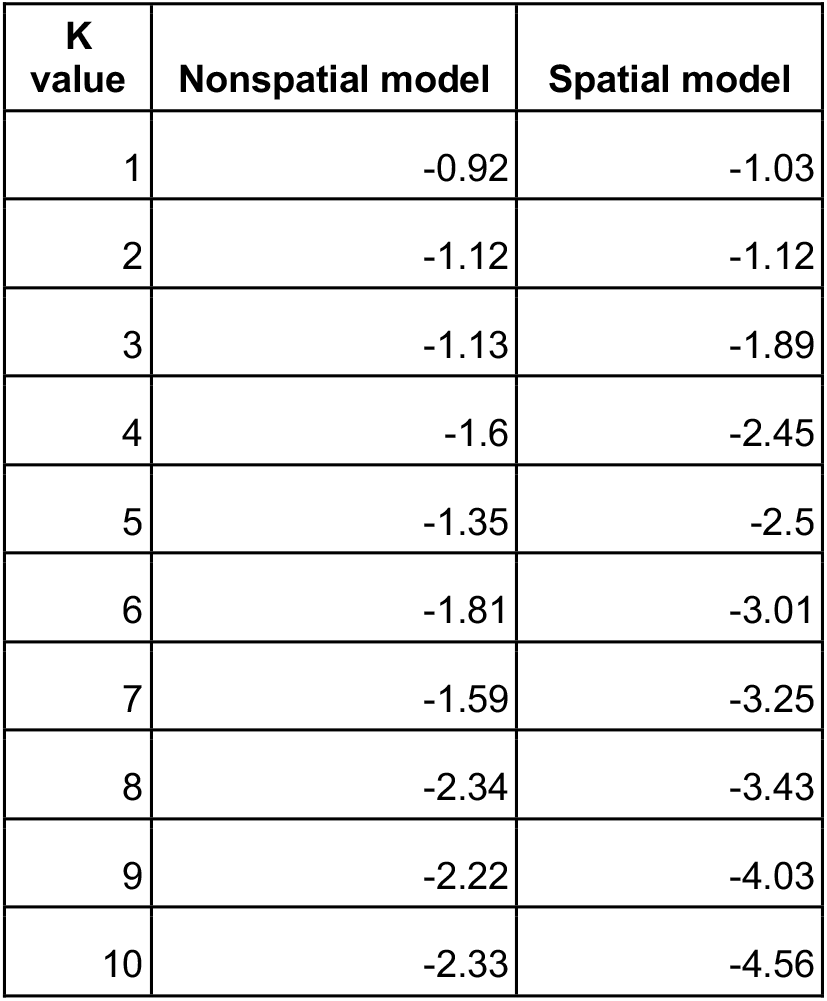
Cross-validation errors for spatial and nonspatial model structure analysis. Cross-validation errors for K values of 1-10 (number of ancestral populations) are shown for spatial and nonspatial models. Cross-validation values closer to zero indicate better model fit. Each model was tested with 1,000 iterations and repeated 10 times. The spatial model incorporates geographic distance between populations, whereas the nonspatial model does not.

| K value | Nonspatial model | Spatial model |
| --- | --- | --- |
| 1 | -0.92 | -1.03 |
| 2 | -1.12 | -1.12 |
| 3 | -1.13 | -1.89 |
| 4 | -1.6 | -2.45 |
| 5 | -1.35 | -2.5 |
| 6 | -1.81 | -3.01 |
| 7 | -1.59 | -3.25 |
| 8 | -2.34 | -3.43 |
| 9 | -2.22 | -4.03 |
| 10 | -2.33 | -4.56 |

**FIGURE S4.**
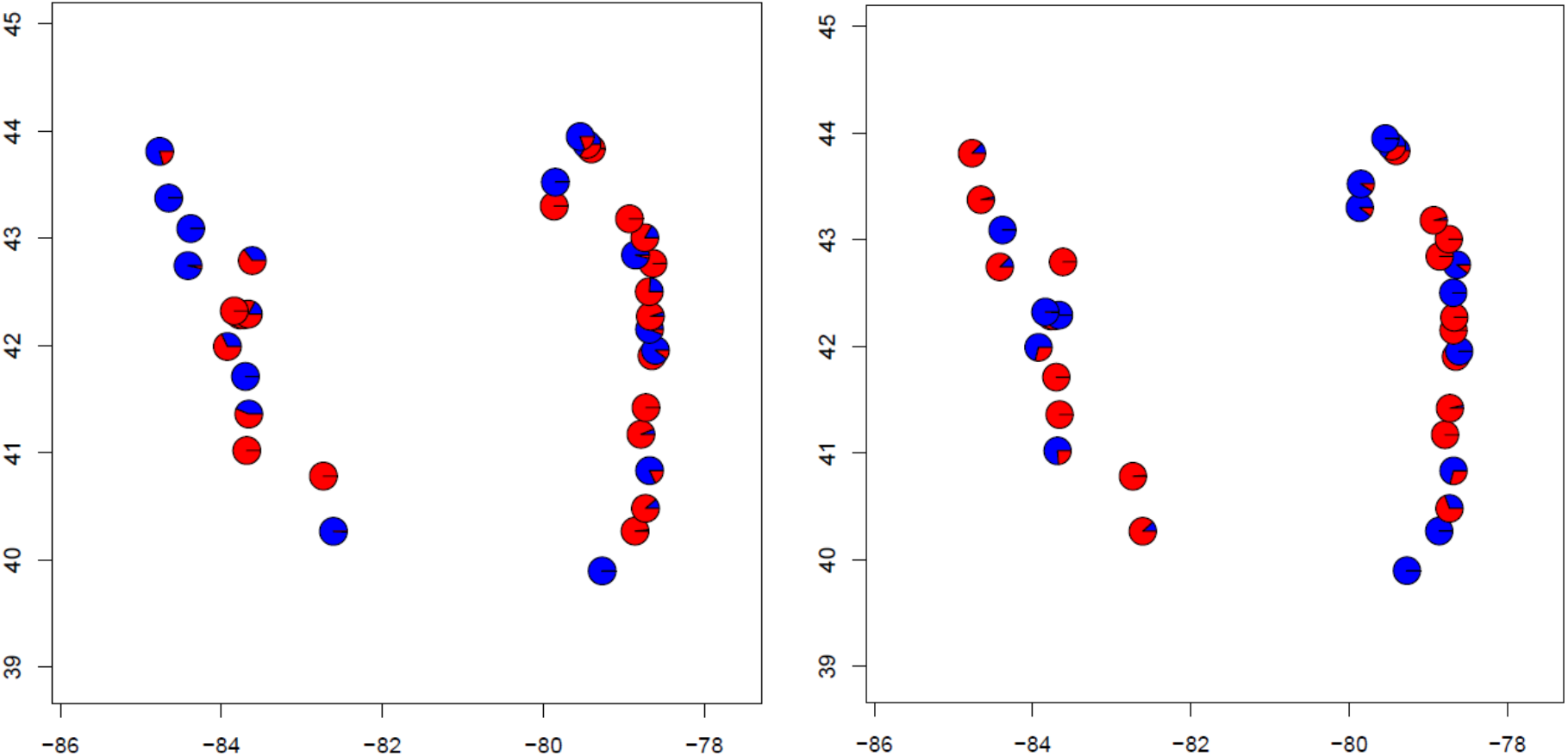
No geographic pattern in population structure at K=2. While K=1 was best supported by structure analysis, results for K=2 do not show clustering of populations by transect or latitude in either the nonspatial (left) or spatial (right) models. This is consistent with an absence of meaningful geographic population structure, as indicated by the strongest support for K=1.

**TABLE S8.**
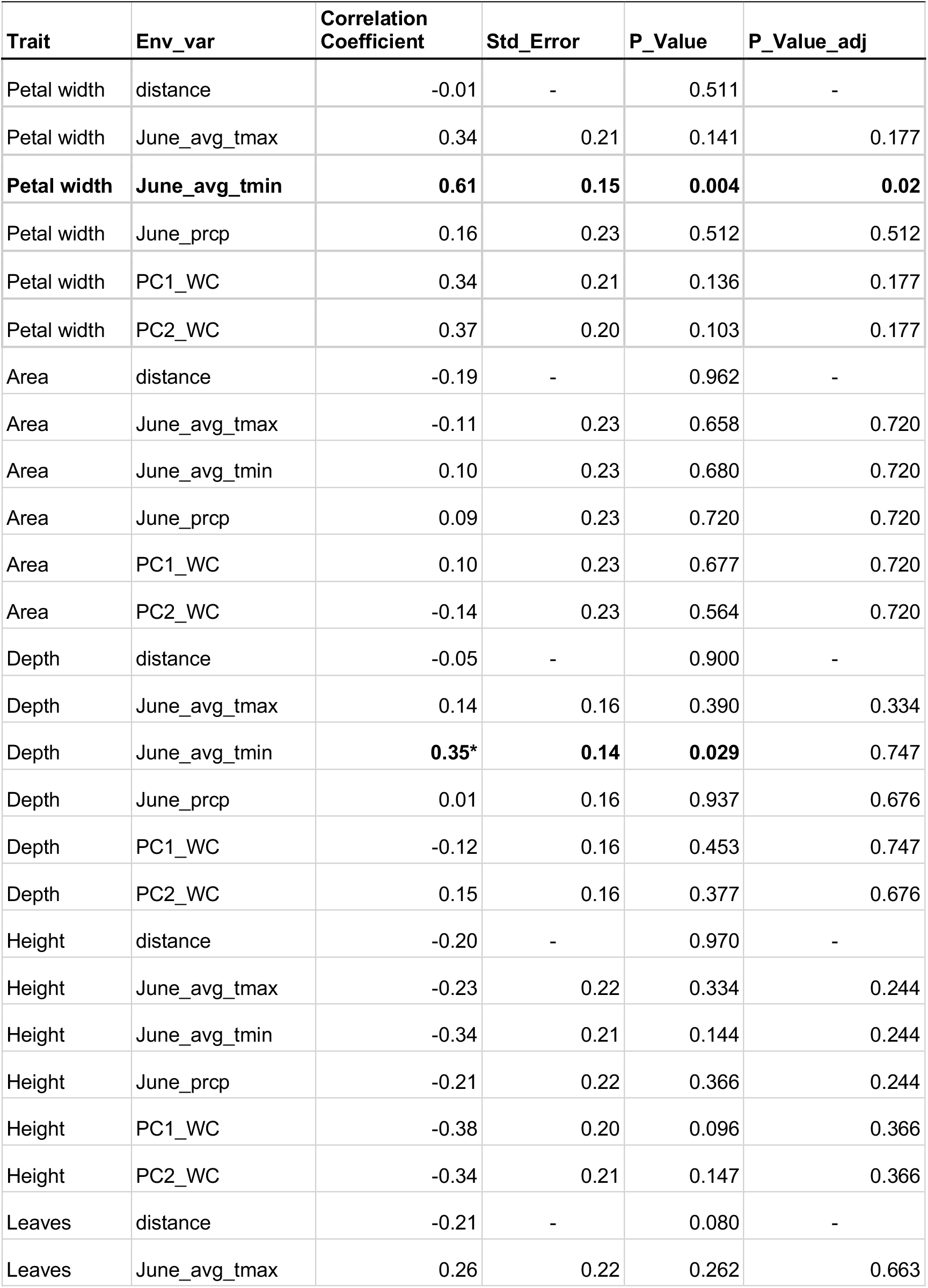
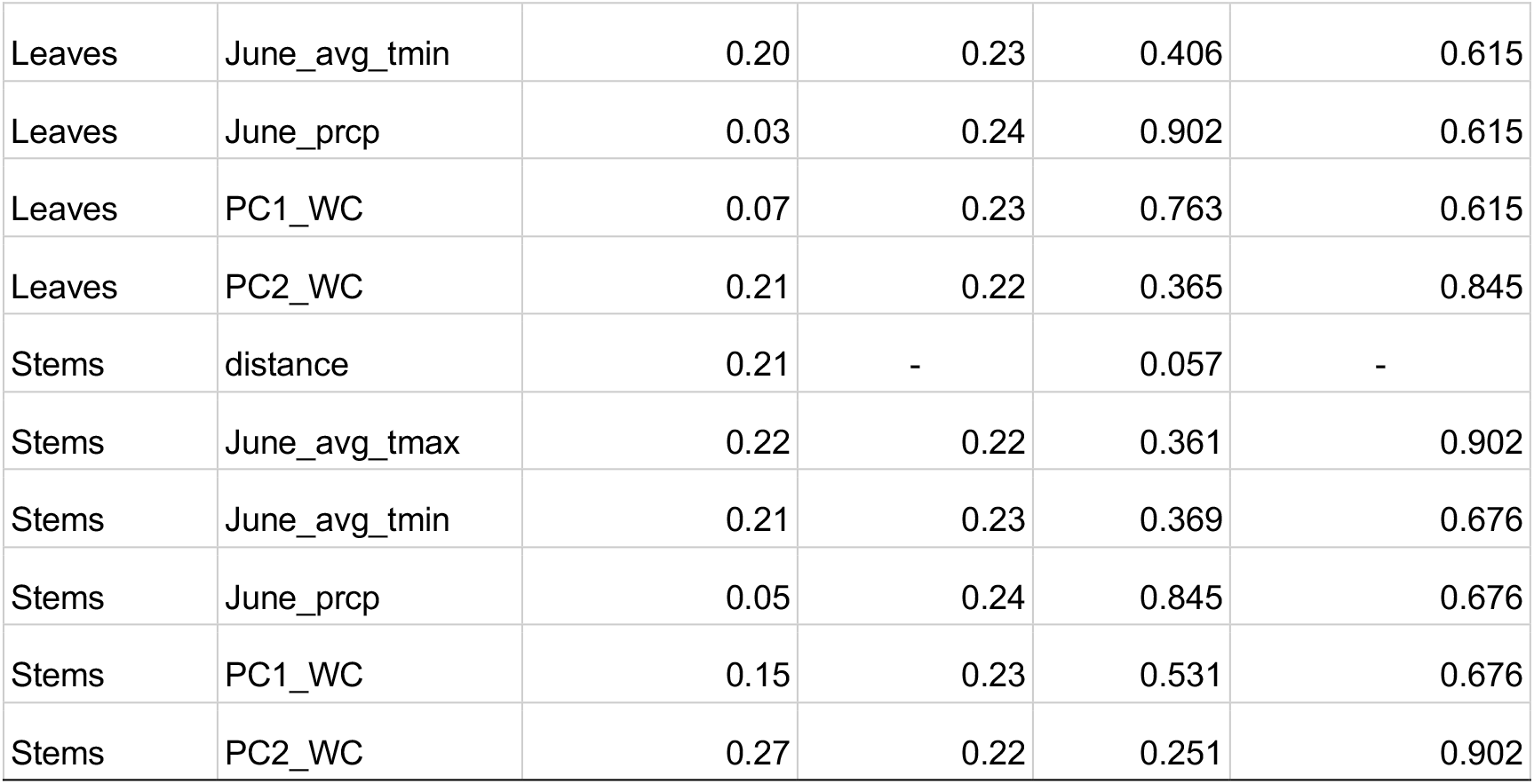
The results of Mantel testing for clinal variation in either geographic distance (Distance) or environmental variables of interest (June_avg_tmax = daily high temperature, June_avg_tmin = June daily low temperature, June_prcp = daily precipitation data, PC1_WC = WorldClim PC1, PC2_WC = WorldClim PC2, see Figure S2).

| Trait | Env_var | Correlation Coefficient | Std_Error | P_Value | P_Value_adj |
| --- | --- | --- | --- | --- | --- |
| Petal width | distance | -0.01 | - | 0.511 | - |
| Petal width | June_avg_tmax | 0.34 | 0.21 | 0.141 | 0.177 |
| **Petal width** | **June_avg_tmin** | **0.61** | **0.15** | **0.004** | **0.02** |
| Petal width | June_prcp | 0.16 | 0.23 | 0.512 | 0.512 |
| Petal width | PC1_WC | 0.34 | 0.21 | 0.136 | 0.177 |
| Petal width | PC2_WC | 0.37 | 0.20 | 0.103 | 0.177 |
| Area | distance | -0.19 | - | 0.962 | - |
| Area | June_avg_tmax | -0.11 | 0.23 | 0.658 | 0.720 |
| Area | June_avg_tmin | 0.10 | 0.23 | 0.680 | 0.720 |
| Area | June_prcp | 0.09 | 0.23 | 0.720 | 0.720 |
| Area | PC1_WC | 0.10 | 0.23 | 0.677 | 0.720 |
| Area | PC2_WC | -0.14 | 0.23 | 0.564 | 0.720 |
| Depth | distance | -0.05 | - | 0.900 | - |
| Depth | June_avg_tmax | 0.14 | 0.16 | 0.390 | 0.334 |
| Depth | June_avg_tmin | **0.35\*** | **0.14** | **0.029** | 0.747 |
| Depth | June_prcp | 0.01 | 0.16 | 0.937 | 0.676 |
| Depth | PC1_WC | -0.12 | 0.16 | 0.453 | 0.747 |
| Depth | PC2_WC | 0.15 | 0.16 | 0.377 | 0.676 |
| Height | distance | -0.20 | - | 0.970 | - |
| Height | June_avg_tmax | -0.23 | 0.22 | 0.334 | 0.244 |
| Height | June_avg_tmin | -0.34 | 0.21 | 0.144 | 0.244 |
| Height | June_prcp | -0.21 | 0.22 | 0.366 | 0.244 |
| Height | PC1_WC | -0.38 | 0.20 | 0.096 | 0.366 |
| Height | PC2_WC | -0.34 | 0.21 | 0.147 | 0.366 |
| Leaves | distance | -0.21 | - | 0.080 | - |
| Leaves | June_avg_tmax | 0.26 | 0.22 | 0.262 | 0.663 |
| Leaves | June_avg_tmin | 0.20 | 0.23 | 0.406 | 0.615 |
| Leaves | June_prdp | 0.03 | 0.24 | 0.902 | 0.615 |
| Leaves | PC1_WC | 0.07 | 0.23 | 0.763 | 0.615 |
| Leaves | PC2_WC | 0.21 | 0.22 | 0.365 | 0.845 |
| Stems | distance | 0.21 | - | 0.057 | - |
| Stems | June_avg_tmax | 0.22 | 0.22 | 0.361 | 0.902 |
| Stems | June_avg_tmin | 0.21 | 0.23 | 0.369 | 0.676 |
| Stems | June_prdp | 0.05 | 0.24 | 0.845 | 0.676 |
| Stems | PC1_WC | 0.15 | 0.23 | 0.531 | 0.676 |
| Stems | PC2_WC | 0.27 | 0.22 | 0.251 | 0.902 |

**TABLE S9.**
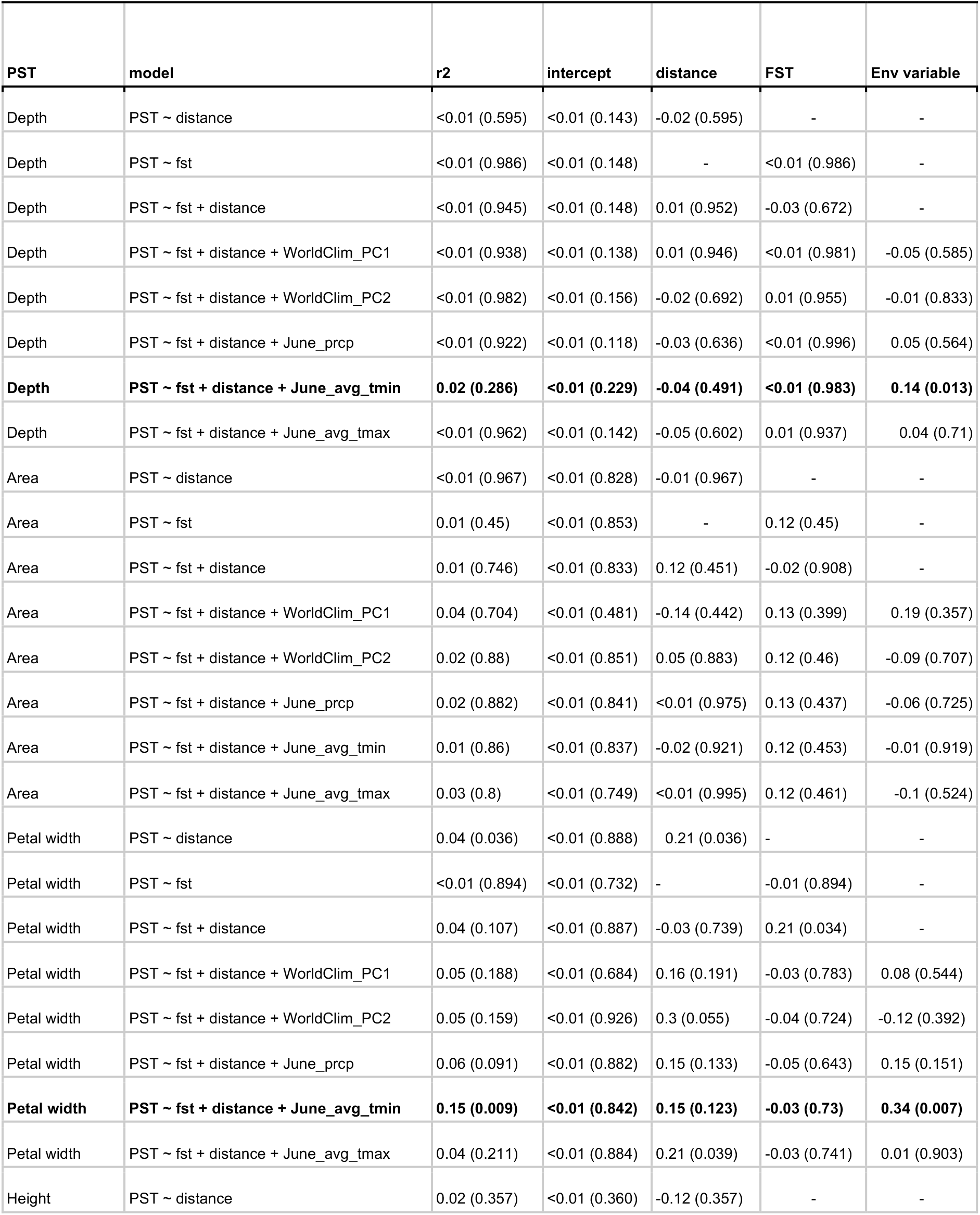
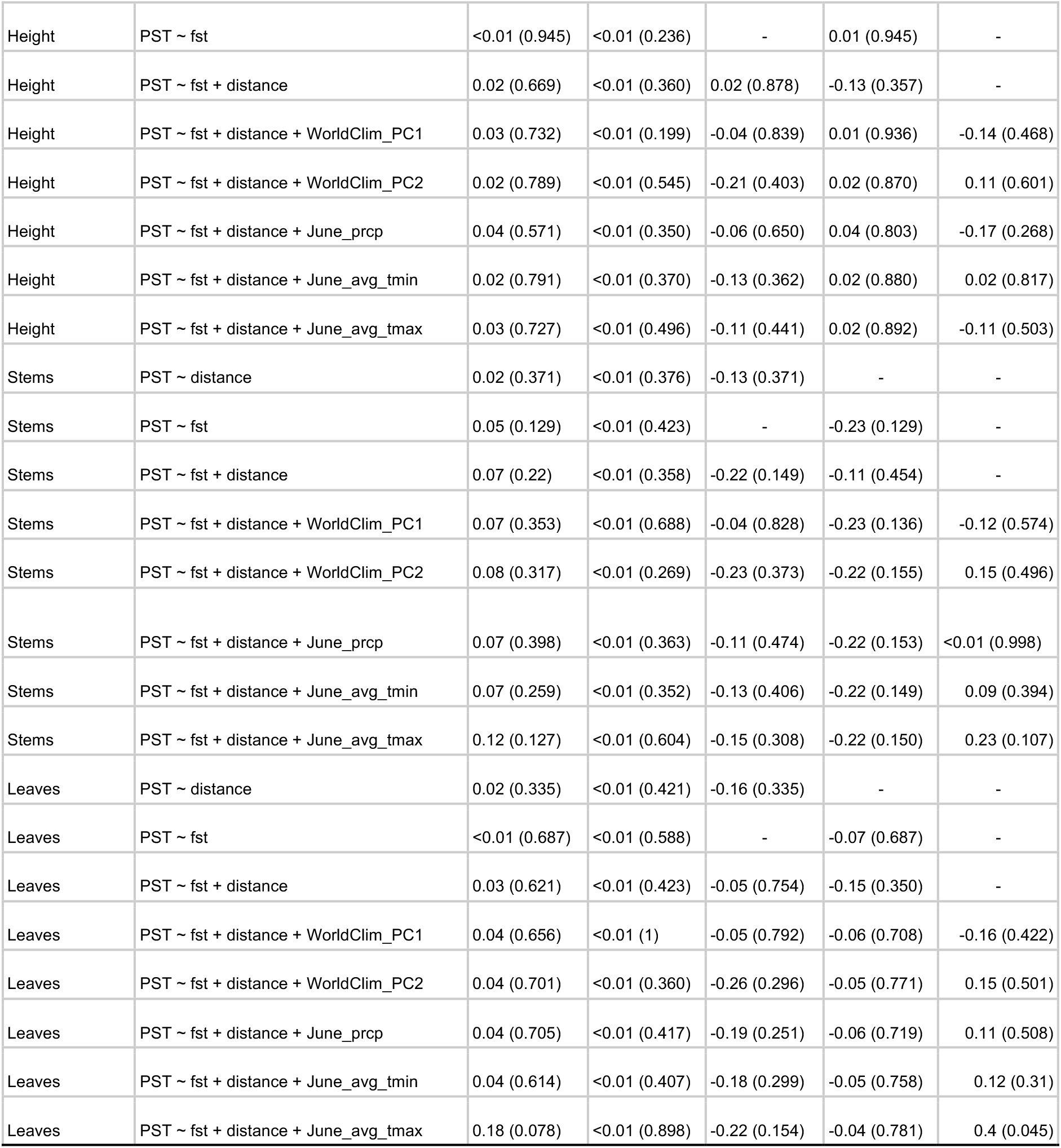
Results of multiple regression of distance matrices (MRM). For each trait (Depth, Area, Petal width, Height, Stems, Leaves), we tested whether P_ST_ was explained by the first 2 WorldClim PC axes or by three June weather variables, while accounting for pairwise geographic distance and F_ST_ between sites. Depth was evaluated across both transects, all other traits were evaluated for Midwest sites only. We report MRM r^2^ values with P-values in parenthesis, and bold those that are significant.

| PST | model | r<sup>2</sup> | intercept | distance | FST | Env variable |
| --- | --- | --- | --- | --- | --- | --- |
| Depth | PST ~ distance | <0.01 (0.595) | <0.01 (0.143) | -0.02 (0.595) | - | - |
| Depth | PST ~ fst | <0.01 (0.986) | <0.01 (0.148) | - | <0.01 (0.986) | - |
| Depth | PST ~ fst + distance | <0.01 (0.945) | <0.01 (0.148) | 0.01 (0.952) | -0.03 (0.672) | - |
| Depth | PST ~ fst + distance + WorldClim_PC1 | <0.01 (0.938) | <0.01 (0.138) | 0.01 (0.946) | <0.01 (0.981) | -0.05 (0.585) |
| Depth | PST ~ fst + distance + WorldClim_PC2 | <0.01 (0.982) | <0.01 (0.156) | -0.02 (0.692) | 0.01 (0.955) | -0.01 (0.833) |
| Depth | PST ~ fst + distance + June_prdp | <0.01 (0.922) | <0.01 (0.118) | -0.03 (0.636) | <0.01 (0.996) | 0.05 (0.564) |
| **Depth** | **PST ~ fst + distance + June_avg_tmin** | **0.02 (0.286)** | **<0.01 (0.229)** | **-0.04 (0.491)** | **<0.01 (0.983)** | **0.14 (0.013)** |
| Depth | PST ~ fst + distance + June_avg_tmax | <0.01 (0.962) | <0.01 (0.142) | -0.05 (0.602) | 0.01 (0.937) | 0.04 (0.71) |
| Area | PST ~ distance | <0.01 (0.967) | <0.01 (0.828) | -0.01 (0.967) | - | - |
| Area | PST ~ fst | 0.01 (0.45) | <0.01 (0.853) | - | 0.12 (0.45) | - |
| Area | PST ~ fst + distance | 0.01 (0.746) | <0.01 (0.833) | 0.12 (0.451) | -0.02 (0.908) | - |
| Area | PST ~ fst + distance + WorldClim_PC1 | 0.04 (0.704) | <0.01 (0.481) | -0.14 (0.442) | 0.13 (0.399) | 0.19 (0.357) |
| Area | PST ~ fst + distance + WorldClim_PC2 | 0.02 (0.88) | <0.01 (0.851) | 0.05 (0.883) | 0.12 (0.46) | -0.09 (0.707) |
| Area | PST ~ fst + distance + June_prdp | 0.02 (0.882) | <0.01 (0.841) | <0.01 (0.975) | 0.13 (0.437) | -0.06 (0.725) |
| Area | PST ~ fst + distance + June_avg_tmin | 0.01 (0.86) | <0.01 (0.837) | -0.02 (0.921) | 0.12 (0.453) | -0.01 (0.919) |
| Area | PST ~ fst + distance + June_avg_tmax | 0.03 (0.8) | <0.01 (0.749) | <0.01 (0.995) | 0.12 (0.461) | -0.1 (0.524) |
| Petal width | PST ~ distance | 0.04 (0.036) | <0.01 (0.888) | 0.21 (0.036) | - | - |
| Petal width | PST ~ fst | <0.01 (0.894) | <0.01 (0.732) | - | -0.01 (0.894) | - |
| Petal width | PST ~ fst + distance | 0.04 (0.107) | <0.01 (0.887) | -0.03 (0.739) | 0.21 (0.034) | - |
| Petal width | PST ~ fst + distance + WorldClim_PC1 | 0.05 (0.188) | <0.01 (0.684) | 0.16 (0.191) | -0.03 (0.783) | 0.08 (0.544) |
| Petal width | PST ~ fst + distance + WorldClim_PC2 | 0.05 (0.159) | <0.01 (0.926) | 0.3 (0.055) | -0.04 (0.724) | -0.12 (0.392) |
| Petal width | PST ~ fst + distance + June_prdp | 0.06 (0.091) | <0.01 (0.882) | 0.15 (0.133) | -0.05 (0.643) | 0.15 (0.151) |
| **Petal width** | **PST ~ fst + distance + June_avg_tmin** | **0.15 (0.009)** | **<0.01 (0.842)** | **0.15 (0.123)** | **-0.03 (0.73)** | **0.34 (0.007)** |
| Petal width | PST ~ fst + distance + June_avg_tmax | 0.04 (0.211) | <0.01 (0.884) | 0.21 (0.039) | -0.03 (0.741) | 0.01 (0.903) |
| Height | PST ~ distance | 0.02 (0.357) | <0.01 (0.360) | -0.12 (0.357) | - | - |
| Height | PST ~ fst | <0.01 (0.945) | <0.01 (0.236) | - | 0.01 (0.945) | - |
| Height | PST ~ fst + distance | 0.02 (0.669) | <0.01 (0.360) | 0.02 (0.878) | -0.13 (0.357) | - |
| Height | PST ~ fst + distance + WorldClim_PC1 | 0.03 (0.732) | <0.01 (0.199) | -0.04 (0.839) | 0.01 (0.936) | -0.14 (0.468) |
| Height | PST ~ fst + distance + WorldClim_PC2 | 0.02 (0.789) | <0.01 (0.545) | -0.21 (0.403) | 0.02 (0.870) | 0.11 (0.601) |
| Height | PST ~ fst + distance + June_prdp | 0.04 (0.571) | <0.01 (0.350) | -0.06 (0.650) | 0.04 (0.803) | -0.17 (0.268) |
| Height | PST ~ fst + distance + June_avg_tmin | 0.02 (0.791) | <0.01 (0.370) | -0.13 (0.362) | 0.02 (0.880) | 0.02 (0.817) |
| Height | PST ~ fst + distance + June_avg_tmax | 0.03 (0.727) | <0.01 (0.496) | -0.11 (0.441) | 0.02 (0.892) | -0.11 (0.503) |
| Stems | PST ~ distance | 0.02 (0.371) | <0.01 (0.376) | -0.13 (0.371) | - | - |
| Stems | PST ~ fst | 0.05 (0.129) | <0.01 (0.423) | - | -0.23 (0.129) | - |
| Stems | PST ~ fst + distance | 0.07 (0.22) | <0.01 (0.358) | -0.22 (0.149) | -0.11 (0.454) | - |
| Stems | PST ~ fst + distance + WorldClim_PC1 | 0.07 (0.353) | <0.01 (0.688) | -0.04 (0.828) | -0.23 (0.136) | -0.12 (0.574) |
| Stems | PST ~ fst + distance + WorldClim_PC2 | 0.08 (0.317) | <0.01 (0.269) | -0.23 (0.373) | -0.22 (0.155) | 0.15 (0.496) |
| Stems | PST ~ fst + distance + June_prdp | 0.07 (0.398) | <0.01 (0.363) | -0.11 (0.474) | -0.22 (0.153) | <0.01 (0.998) |
| Stems | PST ~ fst + distance + June_avg_tmin | 0.07 (0.259) | <0.01 (0.352) | -0.13 (0.406) | -0.22 (0.149) | 0.09 (0.394) |
| Stems | PST ~ fst + distance + June_avg_tmax | 0.12 (0.127) | <0.01 (0.604) | -0.15 (0.308) | -0.22 (0.150) | 0.23 (0.107) |
| Leaves | PST ~ distance | 0.02 (0.335) | <0.01 (0.421) | -0.16 (0.335) | - | - |
| Leaves | PST ~ fst | <0.01 (0.687) | <0.01 (0.588) | - | -0.07 (0.687) | - |
| Leaves | PST ~ fst + distance | 0.03 (0.621) | <0.01 (0.423) | -0.05 (0.754) | -0.15 (0.350) | - |
| Leaves | PST ~ fst + distance + WorldClim_PC1 | 0.04 (0.656) | <0.01 (1) | -0.05 (0.792) | -0.06 (0.708) | -0.16 (0.422) |
| Leaves | PST ~ fst + distance + WorldClim_PC2 | 0.04 (0.701) | <0.01 (0.360) | -0.26 (0.296) | -0.05 (0.771) | 0.15 (0.501) |
| Leaves | PST ~ fst + distance + June_prdp | 0.04 (0.705) | <0.01 (0.417) | -0.19 (0.251) | -0.06 (0.719) | 0.11 (0.508) |
| Leaves | PST ~ fst + distance + June_avg_tmin | 0.04 (0.614) | <0.01 (0.407) | -0.18 (0.299) | -0.05 (0.758) | 0.12 (0.31) |
| Leaves | PST ~ fst + distance + June_avg_tmax | 0.18 (0.078) | <0.01 (0.898) | -0.22 (0.154) | -0.04 (0.781) | 0.4 (0.045) |

**TABLE S10.** Regression analyses showing the linear, quadratic, and correlational selection gradients acting on each flower trait (color, size (area), petal width) and each plant size trait (height, number of stems). Linear values are from a regression with no quadratic or interaction terms included.

| Trait | | $\beta$ (±se) | P |
| --- | --- | --- | --- |
| Height |  | **0.297 (0.040)** | **<0.001** |
| Stem number |  | **0.441 (0.038)** | **<0.001** |
| Area |  | 0.009 (0.057) | 0.869 |
| Petal width |  | 0.053 (0.057) | 0.351 |
| Color |  | 0.075 (0.084) | 0.371 |
| | | $\gamma$ (±se) | P |
| Height x Height |  | 0.041 (0.068) | 0.546 |
| Stem number x Stem number |  | 0.175 (0.053) | **0.001** |
| Area x Area |  | 0.049 (0.065) | 0.452 |
| Petal width x Petal width |  | -0.069 (0.067) | 0.303 |
| Height x Color |  | **-0.273 (0.088)** | **0.002** |
| Stem number x Color |  | **0.492 (0.090)** | **<0.001** |
| Area x Color |  | 0.115 (0.131) | 0.379 |
| Petal width x Color |  | -0.200 (0.134) | 0.136 |
| Petal width x Stem number |  | 0.089 (0.069) | 0.196 |
| Petal width x Height |  | 0.009 (0.065) | 0.887 |
| Stem number x Area |  | -0.004 (0.073) | 0.957 |
| Height x Area |  | 0.026 (0.065) | 0.692 |
| Petal width x Stem number x Color |  | -0.259 (0.151) | 0.087^ |
| Petal width x Height x Color |  | 0.095 (0.134) | 0.479 |
| Stem number x Area x Color |  | 0.025 (0.151) | 0.871 |
| Stem number x Height x Color |  | **-0.205 (0.056)** | **<0.001** |
| Area x Height x Color |  | -0.013 (0.134) | 0.923 |

**FIGURE S5.**
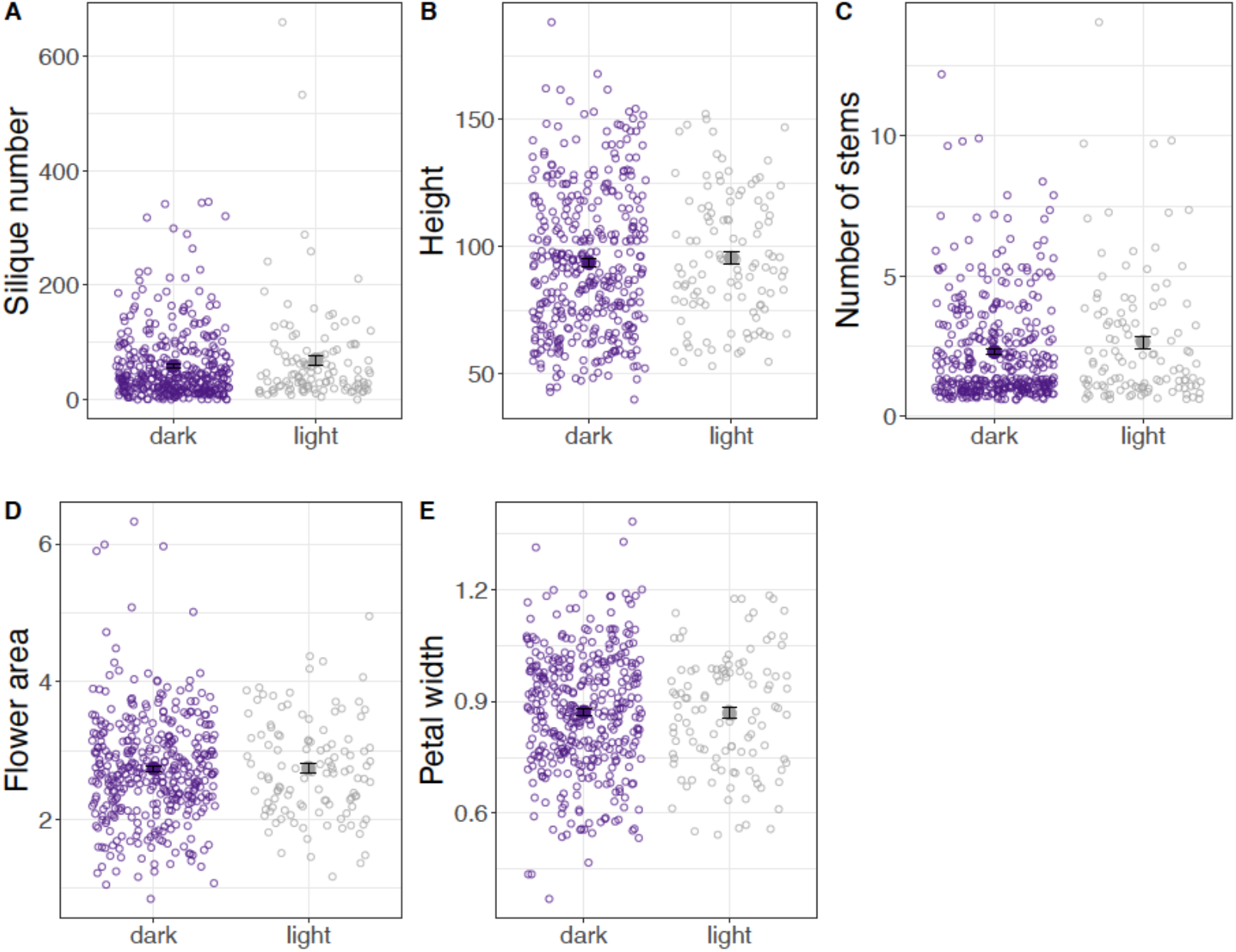
There is no evidence that the traits measured across populations (silique number, plant height, number of stems, and flower area and petal width) differ according to flower color (dark, light).

**TABLE S11.**
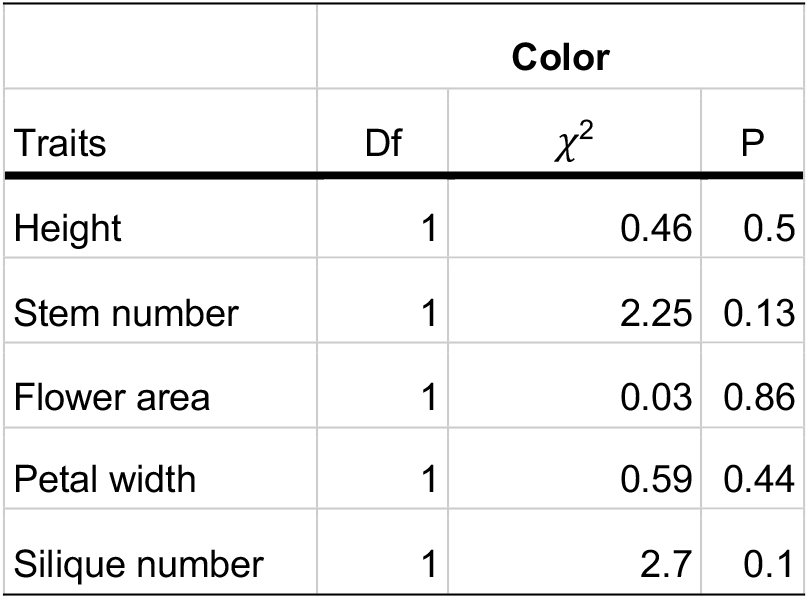
Analyses of variance examining the potential that traits differ according to flower color in *Hesperis matronalis*.

| Traits | Color |  |  |
| --- | --- | --- | --- |
| | Df | $\chi^2$ | P |
| Height | 1 | 0.46 | 0.5 |
| Stem number | 1 | 2.25 | 0.13 |
| Flower area | 1 | 0.03 | 0.86 |
| Petal width | 1 | 0.59 | 0.44 |
| Silique number | 1 | 2.7 | 0.1 |

## Supplemental Methods 2

### Supplemental method 1: ddRADseq Protocol (adapted from Peterson et al. 2012)

Step 1: Extract DNA

Step 2: Quantify & concentrate DNA

Step 3: Anneal adapters P1, P2

Step 4: Double digest

Step 5: Clean digestion

Step 6: Prepare working adapter solution

Step 7: Ligate DNA with adapters

Step 8: Pool samples and clean

Step 9: Size selection

Step 10: PCR

Step 11: Clean & quantify

Step 12: Pool final submission

Step 1: Extract DNA

1. We extracted DNA following the Qiagen plant kit protocol, eluting DNA in 100 uL ultrapure H_2_O.

Step 2: Quantify & concentrate DNA

1. Allow qubit standards to equilibrate to room temperature (from 4°C fridge) for ∼30 min.
2. Label qubit tubes. Label two standards (S1 and S2) and DNA samples (1-N).
3. Take out qubit HS buffer and dye. Gently shake buffer and vortex HS dye to mix.
4. Prepare working solution by making 1:199 dilution of HS dye in buffer in a 15 or 50mL conical tube. I qubit 96 samples together, for which I added 100 uL of Qubit buffer to 19,900 uL Qubit dye (prep for 100 reactions to cover 96 samples, 2 standards, and have extra).
5. Vortex to mix. Place buffer and dye back in drawer after use (dye is light sensitive).
6. Add 190 uL working solution to each standard tube (x2).
7. Add 198 uL working solution to each sample tube (x96).
8. Add 10 uL standard to each standard tube (x2).
9. Add 2 uL DNA sample to each sample tube (x96).
10. Vortex and let sit for 2 min shielded from light. During this time set up table in notebook.
11. On qubit flurometer read standards 1 and 2 for dsDNA HS assay. Then select amount of sample added to each tube (2 uL) and measure each sample. We are aiming for 300ng DNA in 17 uL for our double digest, so at least ∼17.5 ng/uL is the target.
12. For any 100uL samples < 17.5 and > 5 ng/uL that were eluted in ultrapure water, this concentration can be reached using the SpeedVac (low heat, check every 10 minutes and do not evaporate below 20 uL).
13. Qubit any samples from SpeedVac again to ensure > 17.5 ng/uL and quantify for use.

Step 3: Anneal adapters P1, P2

1. We prepared 600 uL of 10x ligation buffer by combining 468 uL H_2_O, 60 uL 5 M NaCl, 60 uL 1 M Tris HCl (ph 8), and 12 uL 0.5 M EDTA.
2. Remove oligos from freezer to thaw. stock adapters from freezer to thaw.
3. Label tubes with adapter combinations. We had 49 tubes - 1 for MspI 2.1/2.2 and 48 for EcoRI 1.1/1.2.
4. We added 40uL Oligo 1 (100uM), 40uL Oligo 2 (100uM), 10uL Ligation Buffer, and 10uL H_2_O for a final adapter concentration of 40uM.
5. On a thermocycler, run a program to heat to 97.5°C for 2.5 minutes. Drop 1 degree every 20 seconds until samples reach room temperature (21°C). Then put on a 4°C hold (run time: ∼30min).

*Annealed adapters can remain at 4°C for a few days, otherwise store at -20°C until use

Step 4: Double digest (∼3 hr)

1. Calculate how much water & DNA is needed for 17ul of DNA for 300 ng of DNA using the DoubleDigest_ConcentrationCalc tab in ddRAD_reaction_tables
  a. Example calculation where [] = DNA concentration from Qubit 300ng/[concentration] = uL DNA to add 17-uL DNA = uL water
2. Pipet appropriate amounts of ultrapure water & then DNA into tubes (17ul). It is less important to have exact amounts of DNA for each individual for this step. You will quantify all individuals again, so approximations can be made [Can take break between adding H_2_O and DNA or after adding DNA. The steps after don’t take too long but you should not pause once you begin to work with the enzymes]
3. In a chilled rack, make a master mix of enzymes and buffer for all tubes. For 96 samples, I find it necessary to prepare the mix for 115 to have enough.
  1. EcoRI HF 0.5ul x number of reactions
  2. Msp 0.5ul x number of reactions
  3. Cutsmart buffer 2ul x number of reactions
4. Place enzymes in freezer immediately after making master mix.
5. Pipet 3ul of the mix into each tube (to get all to 20ul)
6. Vortex & quick spin
7. Run “37digest” on the C1000 Thermocycler-incubates for 3 hours at 37°C, holds at 4°C
8. Store samples in the refrigerator or proceed to section 5 when remove from thermocycler

Step 5: Clean digestion

1. Take aliquot of Ampure Beads out of refrigerator to warm to room temperature (∼30min). If aliquot is not already prepared, allow bottle to warm to room temperature while protected from light and mix well before taking aliquot. 13-14 uL will last for all steps.
2. Prepare 75% EtOH each time cleaning step is performed
  1. Need 400uL/individual
    1. Example Calculation 400uL/individual x 96 individuals = 38400uL = 38.4mL Make 50mL EtOH 0.75 * 50mL = 37.5mL 100% EtOH 50 – 37.5 = 12.5mL diH_2_O
3. Lightly centrifuge PCR strips of DNA
4. Shake Ampure beads well to mix
5. Seal 96 well plate with adhesive film if using new plate.
6. With razor, remove plastic from 2 rows of 96-well plate. Cleaning two strips of DNA at a time is best to avoid letting the beads dry out too much during the process.
7. Pipet 1.5 x (reaction volume) uL of beads into each well of the first 2 rows of the plate. For a 20uL reaction, this is 30uL
8. Using the multi-channel pipet, move all (∼20uL) digested DNA into the 2 rows
  1. Pipet up and down ∼10x to mix thoroughly
  2. Burst any bubbles that may form
9. Let sit at room temperature for 5 minutes
10. Place back on magnetic plate and let sit at room temperature for 3 minutes.
11. Remove and discard all clear liquid using the multi-channel pipet. Do NOT disturb the ring of beads. If you do, repeat the process of 5min and 3min incubations.
  1. You may want to return with a 10 uL multi-channel pipet to get remaining liquid
12. Pour 75% EtOH into plastic trough for easier pipetting
13. Using multichannel pipet, transfer 200uL EtOH into each well
  1. Let sit for 30s
  2. Remove and discard 200 uL EtOH
14. Repeat step 11; return with 10 uL multi-channel pipette to remove all EtOH after wash #2
15. Let sit for 3min or until visibly dry; but do not let the ring dry enough to crack
16. Remove from magnet and add 33uL H_2_O
  1. Pipet up and down ∼10x to mix
  2. May have to scratch at ring with tip to break it up
17. Place back on magnet, let sit for 1min
18. Transfer elutant (∼35uL) to new PCR strips labeled with the corresponding sample names
19. Qubit each cleaned digest to quantify concentration
  1. Repeat Qubit procedure from section 2. You can use 2 uL of each cleaned digest sample for Qubit and each will have at least 33 uL remaining for ligation
  2. Any samples that are <1.75 ng/uL need to be re-digested. If second digest is also low, combine the two, concentrate with SpeedVac, and Qubit that sample again
20. Store in refrigerator

Step 6: Prepare working adapter solution

*We prepared at 10x excess for 1.25 ng/uL DNA and adjusted volume to add to reach 7x excess

**Working solution of annealed MspI is always prepared fresh, I do after step 4 in section 7

1. Dilute 10x ligation (aka annealing) buffer to 1X with ultrapure water.
  1. Should have 50-100 uL extra from section 3 to use here.
2. Determine amount of 40uM annealed stock adapter to add to the ligation conditions using the ligation molarity calculator from Peterson et al. 2012
  1. Edit fields in green for your reaction conditions
    1. Enter your initial DNA mass (concentration after digestion quantification)-this was entered as 1.25 ng/uL DNA
    2. Enter amount of adapter excess (default of 10, anywhere from 5-10 is fine)- we used 7
    3. Enter target adapter volume/reaction (default of 1-2uL is fine)- this varied based on DNA yield and was set so lowest digest had no water added
  2. Calculate amount of each adapter to make
    1. For EcoRI-HF, you will use the same barcode for 2 individuals (48 adapters and 96 individuals in your library). This means if you entered target adapter volume/reaction as “1” uL, you will need at least 2 uL of each diluted adapter but excess can be prepared and frozen for later use.
    2. For MspI, need 1uL per reaction. For 96 samples make 100 uL in order to have enough (don’t make excess for additional libraries, however).
3. Combine correct amounts of buffer and barcodes (EcoRI) in PCR tubes
  1. Using 6, 12-tube PCR strips, combine amount of 1X buffer found in cell B20 and amount of stock annealed adapter found in cell B19 in each of the 48 tubes-for these numbers, I added 1.25 uL of 4x adapter stock to 12.75 uL buffer to make 14 uL of working adapter solution for each EcoRI adapter (x48) in PCR tubes.
  1. Excess can be frozen-as of 11/3/2022 we have some remaining for use
4. Combine correct amounts of buffer (C20) and MspI (C19) in 1.5mL microcentrifuge tube
  a. I added 22.2 MspI stock to 78.8 buffer to make working MspI solution

Step 7: Ligate DNA with adapters

1. Using the DoubleDigest_ConcentrationCalc tab in ddRAD_reaction_tables, enter the post-digest Qubit results to get volume of DNA and water necessary for each sample
  1. Change the formula according to the concentration you want to use. This will be bound by the lowest concentration, and I aimed for 1.75 and used 1.73 ng/uL
  2. Round volumes and sort by rounded DNA or H_2_O volume column. I used volumes to the tenth of a decimal place, but they can be used to the hundredth
2. Label each PCR tube of the 12 strips as 1-96.
3. Pipette ultrapure water into each of the 96 tubes in the volumes from your spreadsheet. By doing the water first, the same pipette tip can be used.
4. Pipette (up to 33 uL) DNA into each of the 96 tubes in the volume from the spreadsheet. Be careful not to contaminate-new pipet tip for reach sample, only open 1 at a time [Now is okay time to take a break, won’t want to pause once thaw materials from -20°C freezer]
5. Prepare 630uL of Msp Master Mix on a chilled rack. MspI adapter in 1x buffer (sec 6) 1ul x number of rxns 105uL 10x ligase buffer 4ul x number of rxns 420uL T4 ligase 1ul x number of rxns 105uL · Return T4 ligase and T4 ligase buffer to -20°C freezer after use
6. Pipet 6uL Master Mix into each tube. Be careful not to contaminate-new pipet tip for reach sample, only open 1 at a time
7. Pipet 1uL of the EcoRI adapters (Set of 48, so you will use each adapter 2x - make sure to keep track of which sample got which adapter set!)
  1. You can use the multi-channel for this
  2. Adapter P1 has 48 different barcodes found in 6 different strip tubes
  1. Add strip 1 to columns 1,7; strip 2 to 2,8; strip 3 to 3,9; strip 4 to 4,10; strip 5 to 5,11; strip 6 to 6;12
8. Vortex & spin tubes down
9. Run “720Ligation” program on C1000 thermocycler: 16°C for 720 minutes [12hrs] 65°C for 10 minutes Cool by decreasing 2°C per 90s until reaching room temperature (21°C) Hold 4°C

Step 8: Pool samples and clean

1. Warm aliquot of Ampure Beads to room temperature protected from light for ∼30 min
2. Will need two, 2ml tubes - one for each unique set of adapters
  1. Combine strips 1-6 into a single tube “A” and strips 7-12 into a second tube “B”
  2. Vortex
3. Split Tube A into 1.5 uL microcentrifuge tubes containing < 400 uL
  1. Sample: For 1920 uL total; use 5 tubes with ∼384 uL in each tube
  2. The last tube will have less, probably between 250-300 uL-It is important to measure this tube and separately calculate how much of the beads will be used.
4. Prepare 15 mL of 75% EtOH if none left from prior cleaning
5. Shake Ampure beads well to mix
6. Pipet 1.5 x (reaction volume) uL of beads into each aliquot from tube “A”
  1. For a 384 uL DNA, this is 576 uL beads
  2. For 300 uL DNA, this is 450 uL
  3. For 250 uL DNA, this is 375 uL
7. Pipet up and down ∼10x to mix thoroughly
8. Let sit at room temperature for 5 minutes
9. Place on magnetic sstand and let sit at room temperature for 5 minutes.
10. Remove and discard all clear liquid. Do NOT disturb the column of beads. If you do, repeat the process of 5min incubations.
  1. Return with a 20 uL pipet to make sure all liquid removed
11. Transfer 650uL 75% EtOH into each well
  1. Let sit for 30s, then remove and discard EtOH
12. Repeat step 11; return with 20 uL pipette to make sure all EtOH is removed after wash #2
13. Let sit for 5 min or until visibly dry; but do not let the column dry enough to crack
14. Remove from magnet and add 40 uL H_2_O
  1. Pipet up and down ∼10x to mix. May have to scratch column with tip to break up
15. Place back on magnet, let sit for 1 min
16. Transfer elutant (∼43uL) to new, single 1.5 mL tube with corresponding sample name. It may help to tilt the top of the 1.5mL tube forward so the bottom is closer to the magnet.
17. Repeat steps 3-16 for tube “B”
18. At this point you will have ∼200 uL sample in each of two tubes, A and B
19. Repeat cleaning procedure again with these two samples.
  1. This time, for 200 uL DNA add 300 uL beads
  2. Mix well with pipette. Make sure to use different tip for each tube
  3. Incubate for 5 min, then place on magnet and incubate for 5 more min
  4. Remove and discard clear liquid
  5. Perform 2 ethanol washes with 500 uL ethanol and let sit for 5 min after wash #2
  6. Remove from magnet, add 30 uL water, and pipette up and down to mix
  7. Place back on magnet, let sit for 1 min
  8. Transfer elutant (∼32uL) to new 1.5 mL tube with corresponding sample name
20. Qubit 1 uL of each sample to quantify-need >33.3 ng/uL (1000 ng DNA) in each tube
21. Store in refrigerator

Step 9: Size selection

1. Bring Pippin prep marker to room temp for ∼30 min
2. Vortex marker
3. Add 10 uL marker L to 30 uL of each DNA sample and vortex to mix well
4. Follow Pippin prep instructions (Jill does this):
  a. Machine will need to be calibrated
  b. Remove adhesive seal from cassette and test for conductivity
  c. Remove 40 uL buffer from elution wells and replace with 40 uL fresh buffer
  d. Run must be set up on computer software
    i. Make sure wells are labeled with sample names
    ii. Make sure unused wells are set as “OF”
2. Add 40 uL sample to each appropriate well
3. Run. Takes ∼90 minutes for 600 bp fragment selection
4. Remove 40 uL from each sample elution well. It can be tricky to recover from bottom of well-make sure to try with small pipette tip to recover all 40 ul!
5. Seal unused cassette wells with adhesive
6. Qubit 1 uL to quantify yield after size selection-should be close to 0.7 ng/uL

Step 10: PCR

1. Take out reagents to thaw from -20 freezer (except Taq-leave in freezer). These will take ∼30 min to thaw. DMSO should be thawed at room temp.
2. Set up two PCR strips for PCR reactions

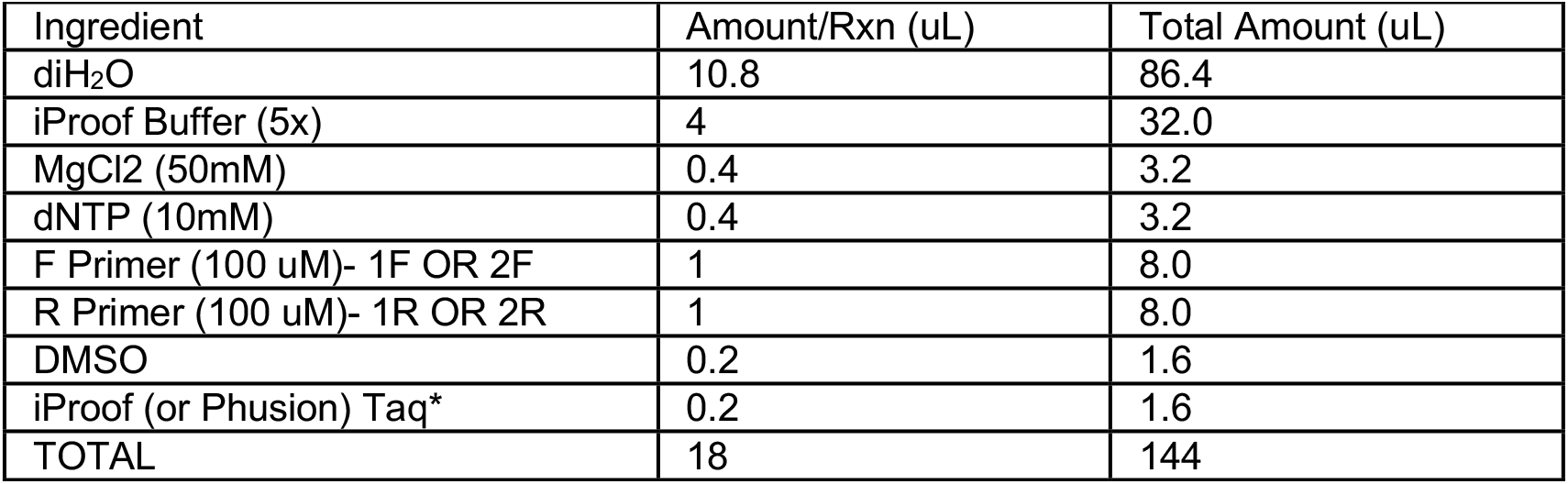
  1. 8 tubes will have Sample A with primers 1.1 and 1.2, 8 will have sample B with primers 2.1 and 2.2
3. On ice, make 2 different master mixes, 1 for each set of primers *Remove Taq from -20 freezer immediately before use and add last
4. Pipette 18 uL master mix to each tube
5. Add 2 uL DNA from library A to first strip of tubes and from library B to second strip
6. Vortex gently and spin down
7. Run “PCR” program in C1000 Thermocycler:
  i. 98°C for 30s
  ii. 98°C for 10s
  iii. 60°C for 20s
  iv. 72°C for 20s Repeat steps ii-iv for 8-12 cycles*
  v. 72°C for 10min
  vi. Hold at 4°C

| Ingredient | Amount/Rxn (uL) | Total Amount (uL) |
| --- | --- | --- |
| diH<sub>2</sub>O | 10.8 | 86.4 |
| iProof Buffer (5x) | 4 | 32.0 |
| MgCl<sub>2</sub> (50mM) | 0.4 | 3.2 |
| dNTP (10mM) | 0.4 | 3.2 |
| F Primer (100 uM)- 1F OR 2F | 1 | 8.0 |
| R Primer (100 uM)- 1R OR 2R | 1 | 8.0 |
| DMSO | 0.2 | 1.6 |
| iProof (or Phusion) Taq* | 0.2 | 1.6 |
| TOTAL | 18 | 144 |

*Number of Cycles: 8 if concentration >= 1.5 ng/ul, 10 if > 0.7 ng/uL, or 12 if < 0.7 ng/ul

Step 11: Clean & quantify

1. Pool across PCR strips
  1. 8 tubes 1-8 for lib A pooled to one 1.5 mL tube with ∼150 uL
  2. 8 tubes 1-8 for lib B pooled to one 1.5 mL tube with ∼150 uL
2. Clean these two samples.
  1. This time, for 150 uL DNA add 225 uL beads
  2. Mix well with pipette. Make sure to use different tip for each tube
  3. Incubate for 5 min, then place on magnet and incubate for 5 more min
  4. Remove and discard clear liquid
  5. Perform 2 ethanol washes with 450 uL ethanol and let dry for 5 min after wash #2
  6. Remove from magnet, add 40 uL water, and pipette up and down to mix
  7. Place back on magnet and let sit for 1 min
  8. Transfer elutant (∼42uL) to new 1.5 mL tube with corresponding sample name
3. Quantify with 2ul for Qubit
  a. Will need at least 10 ng/ul
4. Store samples in fridge

Step 12: Pool final submission

*Must submit 30 uL of one final pooled library with a concentration of at least 10 ng/uL

1. Enter concentrations in “pool” tab of ddRAD_reaction_table spreadsheet to calculate volumes of A and B to add together to get 30 uL of equimolar amounts.
2. Pool specified amounts into the 1.5 mL tube. Store in fridge until submission.

### Supplemental method 2: Floral color measurement with spectrometer

Our set up involved covering the 90-degree hole on the probe holder with a piece of cardboard and constructing a small cardboard platform, with a hole removed where the measurements would take place, to rest the probe holder on during measurements. To set up, we connected the PX-2 light source and flame spectrometer with a 15-pin cable, connected the 6-cable end of the bifurcated Fiber optic cable to the PX-2 light source and the end with 1-cable to the flame spectrometer, and plugged in the PX-2 light source. We then removed the probe cap, placed the probe in the 45-degree slot of the probe holder, and connected the flame spectrometer to the computer.

Before first daily use we made sure the “Flash Mode” setting on the rear of the PX-2 was set to “Single,” turned on the PX-2 light source, and opened the OceanView software. On the Oceanview software, we selected “Spectroscopy Application Wizard” -> “Reflectance,” “Active Acquisition,” and then “Next.” We clicked on add/remove controls and checked the box for “Single Strobe.” Under “Main Controls,” we set “Scans to Average” to 100, “Boxcar width” to “5,” checked the box for “Nonlinearity Correction,” set “Holdoff Time” to 5000, and set “Pulse Width” to “1000.” To warm up the light before first use, under “Single Strobe” we selected “Enabled” to turn on the light source until unchecking the box and selecting “Next” after five minutes.

To calibrate the spectrometer, we placed the probe holder on its cardboard stand with its hold positioned on the hold in the cardboard stand. We placed this complex (probe in its probe holder on cardboard stand) over the white standard and selected “Enabled” under “Single Strube” to turn on the light. We selected Store Reference” and uncheck “Enabled” when finished to turn the light off. We selected “Next,” and repeated this same procedure but on top of the black tabletop and without checking the box for “Strobe/lamp enable” to take a dark standard with as little light as possible and selected “Finish” co complete calibration. This calibration procedure was performed before each population was measured. At this time, we selected “Configure graph saving” to set a directory and file base name for the population to be measured. We set the suffix for filenames as “File Counter” with 2 “Padding Digits,” selected “Save every scan,” and clicked “Apply” and “Exit” when finished.

To measure flower petal reflectance, we placed the probe, in the probe holder with the cardboard platform, on the flower petal so that the opening was over the petal. We checked the box for “Enabled” under the “Single Strobe” section of the “Acquisition Group Window” to start the measurement. Once a preview of the spectra appeared (the probe was adjusted if needed), we selected the save icon and unchecked “Enabled.”

To turn off the system, we made sure all strobe lamp enable options were unchecked, closed the OceanView software, turned off the PX-2 light source, unplugged the PX-2, disconnected the flame spectrometer from the computer, and placed the cap on the probe.

### Supplemental method 3: ImageJ protocol for measurement of flower area and perimeter

Set scale each time open ImageJ

1. Draw straight line along scale 1 cm in length
2. Analyze -> Set Scale; enter length as 1 cm, hit “ok”

Measure area and perimeter of flowers

1. Open the image (File -> open, select appropriate image).
2. Using the freehand selection tool, trace the topmost flower of box #1.
3. Control + M to measure (or Analyze -> Measure). Area and perimeter should show up.
4. Repeat steps 2-3 rest of flowers in box.
5. Repeat steps 2-4 with rest of boxes in image.
6. Make sure to save the file with the area and perimeter data as “Image?_results” before closing, where ? is the image number (first number in image name).
7. Close the image and results file.
8. Repeat steps 1-7 with any additional images.

This same protocol was used to measure flower shape and area for 2022 midwest population flowers. Here, the top left chamber was always measured first, followed by the top right, and the bottom tray chamber was last. With each chamber, flowers were measured in order of top to bottom.

## Notes

### Competing Interest Statement

The authors have declared no competing interest.

### Summary of Updates

The legend for Table 1 was revised; fonts were formatted.

## References

Al Gleason, H., and A. Cronquist. 1964. The natural geography of plants. Columbia University Press, New York, NY.

Andrews, S. 2010. FastQC: a quality control tool for high throughput sequence data.

Bates, D., M. Maechler, and B. Bolker. 2011. lme4: linear mixed-effects models using S4 classes. http://lme4.r-forge.r-project.org/lMMwR/lrgprt.pdf.

Benjamini, Y., and Y. Hochberg. 1995. Controlling the false discovery rate: a practical and powerful approach to multiple testing. Journal of the royal statistical society series b-methodological 57:289–300.

Berardi, A. E., P. D. Fields, J. L. Abbate, and D. R. Taylor. 2016. Elevational divergence and clinal variation in floral color and leaf chemistry in Silene vulgaris. Am. J. Bot. 103:1508–1523. Wiley.

Bradburd, G. S., G. M. Coop, and P. L. Ralph. 2018. Inferring continuous and discrete population genetic structure across space. Genetics 210:33–52.

Brommer, J. E. 2011. Whither Pst? The approximation of Qst by Pst in evolutionary and conservation biology: Whither Pst? J. Evol. Biol. 24:1160–1168. Wiley.

Brooks, M., K. Kristensen, K. van Benthem, A. Magnusson, C. Berg, A. Nielsen, H. Skaug, M. Mächler, and B. Bolker. 2017. GlmmTMB balances speed and flexibility among packages for zero-inflated generalized linear mixed modeling. R J. 9:378. The R Foundation.

Brothers, A. N., and J. W. Atwell. 2014. The role of pollinator-mediated selection in the divergence of floral traits between two closely related plant species. Int. J. Plant Sci. 175:287–295. University of Chicago Press.

Campbell, D. R., N. M. Waser, and M. V. Price. 1996. Mechanisms of Hummingbird-Mediated Selection for Flower width in Ipomopsis aggregata. Ecology 77:1463–1472. Wiley.

Caruso, C. M., K. E. Eisen, R. A. Martin, and N. Sletvold. 2019. A meta-analysis of the agents of selection on floral traits: META-ANALYSIS OF SELECTION ON FLORAL TRAITS. Evolution 73:4–14. Wiley.

Chamberlain, S., and D. Hocking. 2023. rnoaa: “NOAA” Weather Data from R.

Clegg, M. T., and M. Durbin. 2000. Flower color variation: a model for the experimental study of evolution. Proc. Natl. Acad. Sci. U. S. A. 97:7016–7023.

Coberly, L. C., and M. D. Rausher. 2003a. Analysis of a chalcone synthase mutant in Ipomoea purpurea reveals a novel function for flavonoids: amelioration of heat stress. Mol. Ecol. 12:1113–1124. Wiley-Blackwell.

Coberly, L. C., and M. D. Rausher. 2003b. Analysis of a chalcone synthase mutant in Ipomoea purpurea reveals a novel function for flavonoids: amelioration of heat stress. Mol. Ecol. 12:1113–1124. Wiley.

Conner, J. K., and D. L. Hartl. 2004. A primer of ecological genetics. Oxford University Press, New York, NY.

Danecek, P., J. K. Bonfield, J. Liddle, J. Marshall, V. Ohan, M. O. Pollard, A. Whitwham, T. Keane, S. A. McCarthy, R. M. Davies, and H. Li. 2021. Twelve years of SAMtools and BCFtools. Gigascience 10:giab008. Oxford University Press (OUP).

Dellinger, A. S. 2020. Pollination syndromes in the 21st century: where do we stand and where may we go? New Phytol. 228:1193–1213. Wiley.

Delph, L. F., and J. K. Kelly. 2014. On the importance of balancing selection in plants. New Phytol. 201:45–56. Wiley.

Del Valle, J. C., A. Gallardo-López, M. L. Buide, J. B. Whittall, and E. Narbona. 2018. Digital photography provides a fast, reliable, and noninvasive method to estimate anthocyanin pigment concentration in reproductive and vegetative plant tissues. Ecol. Evol. 8:3064–3076. Wiley.

Dray, S., and A.-B. Dufour. 2007. The ade4 Package: Implementing the duality diagram for ecologists. J. Stat. Softw. 22:1–20. Foundation for Open Access Statistic.

Elle, E., and R. Carney. 2003. Reproductive assurance varies with flower size in Collinsia parviflora (Scrophulariaceae). Am. J. Bot. 90:888–896. Wiley.

Epperson, B. K., and M. T. Clegg. 1986. Spatial-Autocorrelation Analysis of Flower Color Polymorphisms within Substructured Populations of Morning Glory (Ipomoea-Purpurea). Am. Nat. 128:840–858. University of Chicago Press.

Fenster, C. B., W. S. Armbruster, P. Wilson, M. R. Dudash, and J. D. Thomson. 2004. Pollination syndromes and floral specialization. Annu. Rev. Ecol. Evol. Syst. 35:375–403. Annual Reviews.

Fox, J., and S. Weisberg. 2018. An R companion to applied regression. 3rd ed. SAGE Publications, Thousand Oaks, CA.

Francis, A., P. B. Cavers, and S. I. Warwick. 2009. The Biology of Canadian Weeds. 140. Hesperis matronalis L. Can. J. Plant Sci. 89:191–206. Canadian Science Publishing.

Frey, F. M. 2004. Opposing natural selection from herbivores and pathogens may maintain floral-color variation in Claytonia virginica (Portulacaceae). Evolution 58:2426–2437. Wiley.

Frey, F. M. 2007. Phenotypic integration and the potential for independent color evolution in a polymorphic spring ephemeral. Am. J. Bot. 94:437–444. Wiley.

Furrer, R., D. Nychka, and S. Sain. 2010. Fields: Tools for Spatial Data.

Galen, C. 1999a. Flowers and enemies: Predation by nectar-thieving ants in relation to variation in floral form of an alpine wildflower, Polemonium viscosum. Oikos 85:426. JSTOR.

Galen, C. 1999b. Why do flowers vary? The functional ecology of variation in flower size and form within natural plant populations. Bioscience 49:631–640.

Galen, C., and J. Cuba. 2001. Down the tube: pollinators, predators, and the evolution of flower shape in the alpine skypilot, Polemonium viscosum. Evolution 55:1963–1971. Oxford University Press (OUP).

Galen, C., R. A. Sherry, and A. B. Carroll. 1999. Are flowers physiological sinks or faucets? Costs and correlates of water use by flowers of Polemonium viscosum. Oecologia 118:461–470. Springer Science and Business Media LLC.

Gervasi, D. D. L., and F. P. Schiestl. 2017. Real-time divergent evolution in plants driven by pollinators. Nat. Commun. 8:14691. Springer Science and Business Media LLC.

Gohil, R. N., and R. Raina. 1987. Polyploidy accompanied by structural alterations in the evolution of Hesperis matronalis L. Cytologia (Tokyo) 52:223–228. International Society of Cytology.

Gómez, J. M. 2003. Herbivory reduces the strength of pollinator-mediated selection in the Mediterranean herb Erysimum mediohispanicum: consequences for plant specialization. Am. Nat. 162:242–256. University of Chicago Press.

Goslee, S., and D. Urban. 2007. The ecodist package for dissimilarity-based analysis of ecological data. Journal of Statistical Software 22:1–19. jstatsoft.org.

Goudet, J. 2005. hierfstat, a package for r to compute and test hierarchical F-statistics. Mol. Ecol. Notes 5:184–186. Wiley.

Harrell, F., Jr. 2024. Hmisc: Harrell Miscellaneous. cran.uib.no.

Hartig, F. 2024. DHARMa: Residual Diagnostics for Hierarchical (Multi-Level / Mixed) Regression Models.

Hedrick, P. W. 2007. Balancing selection. Curr. Biol. 17:R230–1. Elsevier BV.

Holm, S. 1979. A Simple Sequentially Rejective Multiple Test Procedure. Scandinavian Journal of Statistics 6:65–70. [Board of the Foundation of the Scandinavian Journal of Statistics, Wiley].

Irwin, R. E., S. Y. Strauss, S. Storz, A. Emerson, and G. Guibert. 2003. THE ROLE OF HERBIVORES IN THE MAINTENANCE OF A FLOWER COLOR POLYMORPHISM IN WILD RADISH. Ecology 84:1733–1743.

Jombart, T. 2008. adegenet: a R package for the multivariate analysis of genetic markers. Bioinformatics 24:1403–1405. Oxford University Press (OUP).

Jombart, T., and I. Ahmed. 2011. adegenet 1.3-1: new tools for the analysis of genome-wide SNP data. Bioinformatics 27:3070–3071. Oxford University Press (OUP).

Kamvar, Z. N., J. F. Tabima, and N. J. Grünwald. 2014. Poppr: an R package for genetic analysis of populations with clonal, partially clonal, and/or sexual reproduction. PeerJ 2:e281. PeerJ.

Kay, Q. Q. N. 1978. The role of preferential and assortative pollination in the maintenance of flower color polymorphisms. Pages 175–190 in. in A. J. Richards, ed. The pollination of flowers by insects. Academic Press, London, UK.

Klecka, J., J. Hadrava, and P. Koloušková. 2018. Vertical stratification of plant-pollinator interactions in a temperate grassland. PeerJ 6:e4998.

Koski, M. H. 2023. Pollinators exert selection on floral traits in a pollen-limited, narrowly endemic spring ephemeral. Am. J. Bot. 110:e16101. Wiley.

Lande, R., and S. J. Arnold. 1983. The measurement of selection on correlated characters. Evolution 37:1210–1226. Wiley.

Leinonen, T., R. B. O’Hara, J. M. Cano, and J. Merilä. 2008. Comparative studies of quantitative trait and neutral marker divergence: a meta-analysis. J. Evol. Biol. 21:1–17. academic.oup.com.

Li, H. 2013. Aligning sequence reads, clone sequences and assembly contigs with BWA-MEM.

Maad, J., W. S. Armbruster, and C. B. Fenster. 2013. Floral size variation in Campanula rotundifolia (Campanulaceae) along altitudinal gradients: patterns and possible selective mechanisms. Nord. J. Bot. 31:361–371. Wiley.

Majetic, C. J. 2008a. Floral scent variation in Hesperis matronalis (Brassicaceae): assessing potential causes of within- and among-population variation. University of Pittsburgh, Pittsburgh.

Majetic, C. J. 2008b. The evolutionary ecology of floral scent in Hesperis matronalis: assessing the potential for pollinator-mediated natural selection. University of Pittsburgh.

Majetic, C. J., R. A. Raguso, and T.-L. Ashman. 2009. The sweet smell of success: floral scent affects pollinator attraction and seed fitness in Hesperis matronalis. Funct. Ecol. 23:480–487. Wiley.

Majetic, C. J., R. A. Raguso, S. J. Tonsor, and T.-L. Ashman. 2007. Flower color-flower scent associations in polymorphic Hesperis matronalis (Brassicaceae). Phytochemistry 68:865–874. Elsevier BV.

Marshall, M. M., L. C. Batten, D. L. Remington, and E. P. Lacey. 2019. Natural selection contributes to geographic patterns of thermal plasticity in Plantago lanceolata. Ecol. Evol. 9:2945–2963. Wiley.

Martin, M. 2011. Cutadapt removes adapter sequences from high-throughput sequencing reads. EMBnet J. 17:10. EMBnet Stichting.

Merilä, J., and P. Crnokrak. 2001. Comparison of genetic differentiation at marker loci and quantitative traits: Natural selection and genetic differentiation. J. Evol. Biol. 14:892–903. Wiley.

Milano, E. R., A. M. Kenney, and T. E. Juenger. 2016. Adaptive differentiation in floral traits in the presence of high gene flow in scarlet gilia (Ipomopsis aggregata). Mol. Ecol. 25:5862–5875.

Mitchell, R. J., and D. P. Ankeny. 2001. Effects of Local Conspecific Density on Reproductive Success in Penstemon digitalis and Hesperis matronalis. Ohio J. Sci. 101:22–27.

Morente-López, J., J. F. Scheepens, C. Lara-Romero, R. Ruiz-Checa, P. Tabarés, and J. M. Iriondo. 2020. Past selection shaped phenological differentiation among populations at contrasting elevations in a Mediterranean alpine plant. Environ. Exp. Bot. 170:103894. Elsevier BV.

Nikolov, L. A. 2019. Brassicaceae flowers: diversity amid uniformity. J. Exp. Bot. 70:2623–2635. Oxford University Press (OUP).

Oksanen, F. J., G. Blanchet, M. Friendly, R. Kindt, P. Legendre, D. McGlinn, P. R. Minchin, R. B. O’Hara, G. L. Simpson, P. Solymos, and Others. 2017. vegan: Community Ecology Package. R package version 2.4-4. https.CRAN.R-project.org/package=vegan.

Parachnowitsch, A. L., and A. Kessler. 2010. Pollinators exert natural selection on flower size and floral display in Penstemon digitalis. New Phytol. 188:393–402. Wiley.

Pembleton, L. W., N. O. I. Cogan, and J. W. Forster. 2013. StAMPP: an R package for calculation of genetic differentiation and structure of mixed-ploidy level populations. Mol. Ecol. Resour. 13:946– 952. Wiley.

Peterson, R. 2021a. Finding optimal normalizing transformations via bestNormalize. R J. 13:310. The R Foundation.

Peterson, R. A. 2021b. Finding Optimal Normalizing Transformations via bestNormalize.

Plendl, M., G. E. van der Voort, and J. K. Janes. 2024. Assessing floral trait variation in Platanthera dilatata (Orchidaceae) across an elevational gradient. Discov. Plants 1:63. Springer Science and Business Media LLC.

Posit team. 2023. RStudio: Integrated Development Environment for R. Posit Software, PBC, Boston, MA.

Rausher, M. D. 2008. Evolutionary transitions in floral color. Int. J. Plant Sci. 169:7–21. University of Chicago Press.

Rausher, M. D., and J. D. Fry. 1993. Effects of a Locus Affecting Floral Pigmentation in Ipomoea-Purpurea on Female Fitness Components. Genetics 134:1237–1247.

R Core Team. 2023. R: a language and environment for statistical computing.

Russell, L. n.d. Emmeans: estimated marginal means, aka least-squares means. R package version.

Sarkissian, T. S., and L. D. Harder. 2001. Direct and indirect responses to selection on pollen size in Brassica rapa L. J. Evol. Biol. 14:456–468. Wiley.

Schemske, D. W., and P. Bierzychudek. 2001. Perspective: Evolution of flower color in the desert annual Linanthus parryae: Wright revisited. Evolution 55:1269–1282. Wiley.

Schemske, D. W., and P. Bierzychudek. 2007. Spatial differentiation for flower color in the desert annual Linanthus parryae: was Wright right? Evolution 61:2528–2543. Wiley.

Silva, S., and A. Silva. 2018. Pstat: An R package to assess population differentiation in phenotypic traits. R J. 10:447. The R Foundation.

Sokal, R. R., and F. J. Rohlf. 1995. Biometry. 3rd ed. W.H. Freeman, New York, NY.

Stanton, M. L., A. A. Snow, S. N. Handel, and J. Bereczky. 1989. The impact of a flower-color polymorphism on mating patterns in experimental populations of wild radish (Raphanus raphanistrum L.). Evolution 43:335. Oxford University Press (OUP).

Storz, J. F. 2002. Contrasting patterns of divergence in quantitative traits and neutral DNA markers: analysis of clinal variation: GENETIC AND PHENOTYPIC DIVERGENCE. Mol. Ecol. 11:2537– 2551. Wiley.

Strauss, S. Y., and J. B. Whittall. 2006. Non-pollinator agents of selection on floral traits. Ecology and evolution of flowers 120–138.

Streisfeld, M. A., and J. R. Kohn. 2005. Contrasting patterns of floral and molecular variation across a cline in Mimulus aurantiacus. Evolution 59:2548–2559. Wiley.

Subramaniam, B., and M. Rausher. 2000. BALANCING SELECTION ON A FLORAL POLYMORPHISM. Evolution 54:691–695. Blackwell Publishing Inc.

Vaidya, P., A. McDurmon, E. Mattoon, M. Keefe, L. Carley, C.-R. Lee, R. Bingham, and J. T. Anderson. 2018. Ecological causes and consequences of flower color polymorphism in a self-pollinating plant (Boechera stricta). New Phytol. 218:380–392.

Weiß, C. L., M. Pais, L. M. Cano, S. Kamoun, and H. A. Burbano. 2018. nQuire: a statistical framework for ploidy estimation using next generation sequencing. BMC Bioinformatics 19:122.

Whitlock, M. C. 2008. Evolutionary inference fromQST: Evolutionary inference fromQST. Mol. Ecol. 17:1885–1896. Wiley.

Worley, A., A. M. Baker, J. Thompson, and S. Barrett. 2000. Floral display in narcissus: Variation in flower size and number at the species, population, and individual levels. INT. J. PLANT SCI. 161:69–79.

Zu, P., W. U. Blanckenhorn, and F. P. Schiestl. 2016. Heritability of floral volatiles and pleiotropic responses to artificial selection in Brassica rapa. New Phytol. 209:1208–1219. Wiley.

2019. Picard toolkit. Broad Institute.

